# New isolates refine the ecophysiology of the Roseobacter CHAB-I-5 lineage

**DOI:** 10.1101/2024.05.28.596239

**Authors:** V. Celeste Lanclos, Xiaoyuan Feng, Chuankai Cheng, Mingyu Yang, Cole J. Hider, Jordan T. Coelho, Conner Y. Kojima, Shelby J. Barnes, Catie S. Cleveland, Mei Xie, Yanlin Zhao, Haiwei Luo, J. Cameron Thrash

**Affiliations:** Department of Biological Sciences, University of Southern California, Los Angeles, CA, USA; Simon F. S. Li Marine Science Laboratory, School of Life Sciences and State Key Laboratory of Agrobiotechnology, The Chinese University of Hong Kong, Shatin, SAR, Hong Kong, China; Shenzhen Research Institute, The Chinese University of Hong Kong, Shenzhen, China; Fujian Provincial Key Laboratory of Agroecological Processing and Safety Monitoring, College of Life Sciences, Fujian Agriculture and Forestry University, Fuzhou, China

## Abstract

The CHAB-I-5 cluster is a pelagic lineage that can comprise a significant proportion of all roseobacters in surface oceans and have predicted roles in biogeochemical cycling via heterotrophy, aerobic anoxygenic photosynthesis (AAnP), CO oxidation, DMSP degradation, and other metabolisms. Though cultures of CHAB-I-5 have been reported, none have been explored and the best known representative, strain SB2, was lost from culture after obtaining the genome sequence. We have isolated two new CHAB-I-5 representatives, strains US3C007 and FZCC0083, and assembled complete, circularized genomes with 98.7% and 92.5% average nucleotide identities with the SB2 genome. Comparison of these three with 49 other unique CHAB-I-5 metagenome-assembled and single-cell genomes indicated that the cluster represents a genus with two species, and we identified subtle differences in genomic content between the two species subclusters. Metagenomic recruitment from over fourteen hundred samples expanded their known global distribution and highlighted both isolated strains as representative members of the clade. FZCC0083 grew over twice as fast as US3C007 and over a wider range of temperatures. The axenic culture of US3C007 occurs as pleomorphic cells with most exhibiting a coccobacillus/vibrioid shape. We propose the name *Thalassovivens spotae*, gen nov., sp. nov. for the type strain US3C007^T^.

## Introduction

The Roseobacter group is one of the most ecologically successful groups of bacteria found across marine habitats and are often associated with phytoplankton blooms [1–4]. Members of this clade exist as free-living, attached, and in symbiont forms [1] and can make up to 20% of bacteria in coastal regimes [5]. The most abundant roseobacters in the open ocean belong to the Pelagic Roseobacter Cluster (PRC), which are polyphyletic in the Roseobacter phylogenomic tree, but form a cluster in the dendrogram inferred from genome content similarity [3, 6, 7]. This results from multiple Roseobacter lineages that have evolved gene content that is adaptive for nutrient-poor pelagic waters, such as carbon monoxide and inorganic sulfur oxidation, use of dimethylsulfoniopropionate (DMSP) via multiple pathways, a reduction of metal import systems, and a high proportion of ABC transporters, some of which distinguish them from copiotrophic roseobacters [6, 8, 9]. While many Roseobacter species are easily cultured, the PRC contains multiple clusters without currently isolated representatives, including the CHAB-I-5 lineage. Representatives from the CHAB-I-5 cluster have been cultured on multiple occasions but lost [7, 10, 11], for example, strain SB2 was the first [7].

The CHAB-I-5 cluster comprises free-living marine bacteria distributed from tropical to polar latitudes [7, 12] and is one of the most abundant types of Roseobacter in global oceans. It is found in highest abundances near coastal North America and Europe [12] and constituted up to 20% of microbial clones in the Sargasso Sea [1, 13]. In a study of Chesapeake Bay, CHAB-I-5 was the only Roseobacter that did not decrease in abundance along a salinity gradient and was present in samples across salinities from 13.9-30.5 [14]. While some other members of the Roseobacter group typically associate with phytoplankton blooms, this pattern does not seem to hold for CHAB-I-5 [7]. The abundance and distribution of CHAB-I-5 in global ocean waters corresponds to a high activity level in the cluster [1, 7, 12, 14, 15]. Furthermore, CHAB-I-5 phage are abundant in global waters, particularly in the polar and estuarine systems [10]. This abundance, activity, and widespread phage distribution indicate this group is essential to global nutrient cycling, though the mechanisms of these dynamics are still unexplored.

Current predictions of CHAB-I-5 metabolism come from only four partial genomes [7, 12]. CHAB-I-5 appears to be motile with metabolic pathways for aerobic anoxygenic photosynthesis, carbon monoxide oxidation, inorganic sulfur oxidation, DMSP degradation, phosphonate metabolism, and evidence for thiamin and biotin auxotrophy similar to other PRC members [7, 9, 12]. Incomplete genomes have made it unclear whether CHAB-I-5 can use nitrate, nitrite, or reduce sulfur [7]. Furthermore, we have no knowledge of cell volumes, growth rates, or other fundamental physiological characteristics of this group. No CHAB-I-5 isolate has been maintained in culture long enough for experimental analysis except for the recent isolate FZCC0083, which remains uncharacterized except for use in phage isolations [10]. Thus, our current knowledge of CHAB-I-5 remains limited.

Here we present another new strain, US3C007, an axenic representative of the CHAB-I-5 cluster that is readily propagated on artificial seawater medium and reliably revived from frozen stocks. We conducted the first physiological characterization of CHAB-I-5, and the most extensive genomic analysis of the group to date using new, complete genomes from both US3C007 and FZCC0083 and other publicly-available data. We showed the first morphology of a CHAB-I-5 member and examined the growth dynamics of both strains across ranges of salinity and temperature. Additionally, we analyzed the ecological distribution of CHAB-I-5 from an expanded set of global metagenomic samples that span a wide range of marine and estuarine locations. Together, these data constitute the most in-depth investigation of CHAB-I-5 thus far and provide new insights on the genomics and physiology of these organisms.

## Materials and Methods

### US3C007 isolation

We obtained surface water (2m) from the San Pedro Ocean Time series (SPOTs) monthly cruise on 09/16/2020 via CTD cast. The seawater was transported into the lab and filtered through a 2.7µm GF/D filter, stained with 1x Sybr green (Lonza) for 30 minutes in the dark, and cell density was enumerated on a Guava Easy Cyte 5HT flow cytometer (Millipore, Massachusetts, USA) with settings as described previously [16]. We diluted cells to a final concentration of 1 cell/µL in 10mL of sterilized AMS1 artificial seawater medium [17] and inoculated 3 µL of the diluted cell solution into each well of a 96 x 2.1mL well PTFE plate (Radleys, Essex, UK) containing 1.5mL of AMS1 for a final theoretical concentration of 3 cells/well. Plates incubated in the dark without shaking for 2.5 weeks and enumerated as described above. Positive wells (>10^4^ cells/mL) were transferred to Nalgene Oak Ridge PTFE centrifuge tubes (Thermo Fisher, Massachusetts, USA) containing MWH1 medium [18] in an attempt to move the cultures to a more frequently used medium for convenience. Subsequent transfers of isolates in MWH1 were not successful, so we transferred the initial cultures in the Oak Ridge tubes containing MWH1 to acid-washed 125 ml polycarbonate flasks containing the original isolation medium, AMS1, and growth resumed. The culture has been maintained in this manner over continual transfers. Cultures were cryopreserved in both 10% DMSO and 10% glycerol diluted with AMS1. We grew US3C007 to late-log phase and filtered the cells onto a 0.2µm polycarbonate filter (Millipore) and extracted its DNA using a GenElute Bacterial Genomic DNA Kit (Sigma-Aldrich Co, Darmstadt, Germany). We amplified the DNA and purified the PCR products as previously reported [19], and sent samples for Sanger sequencing to Genewiz (Azenta Life Sciences, New Jersey, USA). We inspected the resulting chromatograms to verify purity through a lack of multiple peaks for a given base call, assembled a contiguous sequence from the forward and reverse complement sequences using CAP3 (https://doua.prabi.fr/software/cap3), and used the web-based NCBI BLASTn with the nr/nt database for sequence identification.

### 16S rRNA gene phylogeny to determine placement within CHAB-I-5

We created a 16S rRNA gene phylogeny to verify placement of US3C007 within the CHAB-I-5 cluster using the Alphaproteobacteria tree and methods from previous work [18, 19] with the addition of known CHAB-I-5 relatives including SB2 [7], three CHAB-I-5 SAGs [12], the original CHAB-I-5 clone [20], and US3C007. We aligned sequences with muscle v3.8.1551 [21], trimmed with trimal v1.4.1 [22], and inferred the phylogeny with IQ-TREE v2.0.6 with flag “-B 1000” [23]. The phylogeny was visualized with Figtree v1.4.4 and all nodes were collapsed except for the branches containing CHAB-I-5 and PRC member HIMB11 to highlight US3C007’s inclusion within the CHAB-I-5 (**Fig. S1**).

### Genome sequencing and assembly

We revived US3C007 from cryostocks and grew the culture in multiple 1L batches to gather DNA for genome sequencing. We filtered the cells onto 0.1µm polycarbonate filters (Millipore) and extracted DNA with a phenol chloroform approach (https://www.protocols.io/view/modified-phenol-chloroform-genomic-dna-extraction-e6nvwkjzwvmk/v2). DNA was pooled together and sent for Illuminia NextSeq 2000 paired end (2×151bp) sequencing at the Microbial Genome Sequencing Center (MiGS) (Pittsburgh, Pennsylvania, USA). Illumina libraries were prepared with the Illumina DNA Prep kit and 10bp UDI indices. Demultiplexing, quality control and adapter trimming was performed with bcl2fastq (v2.20.0422) (https://support.illumina.com/sequencing/sequencing_software/bcl2fastq-conversion-software.html). Illumina reads were trimmed using Trimmomatic (v0.38) to remove poor quality bases [24]. We also performed long-read sequencing in-house using an Oxford Nanopore MinION with a R9.4.1 (FLOMIN106) flow cell (Oxford, UK). For Nanopore sequencing, DNA was sheared with a size selection of 20,000bp or greater using Covaris g-tubes (D-Mark Biosystems, Woburn, USA) and we constructed libraries with the SQK-LSK108 genomic DNA ligation kit (Oxford Nanopore, UK) with modifications (https://doi.org/10.17504/protocols.io.bixskfne). Reads were base-called with Guppy v4.4.1 [25], and demultiplexed using Porechop v0.2.4 (https://github.com/rrwick/Porechop). We assembled the long-read sequence data using Flye v2.9.1 [26] using the “nano-hq” setting and 4 rounds of polishing with minimap [27], included in the Flye assembler. We then used short-reads from Illumina to further improve the assembly with Polypolish v0.5.0 [28]. The resulting assembly was visualized for completion with Bandage v0.8.1 [29].

Bacterial cultivation and DNA extraction of FZCC0083 were performed following our previous paper [10]. Briefly, a surface water sample was collected from the coast of the East China Sea. The FZCC0083 strain was isolated following the dilution cultivation procedure [11], and genomic DNA was extracted using EZ.N.A. Library preparation and genome sequencing was performed following the standard protocols for Illumina sequencing on BGISEQ500 platform (PE100, Qingdao Huada Gene Biotechnology Co., Ltd) [30] and Nanopore sequencing on a Nanopore MinION sequencer (Oxford Nanopore Technologies Inc.) with a R9.4.1 (FLO-MIN106D) flow cell and the SQK-LSK109 genomic DNA ligation kit (Oxford Nanopore, UK). The Illumina sequencing reads (coverage > 200x) were quality trimmed using Trimmomatic v0.36 [24] with options ‘SLIDINGWINDOW:4:15 MAXINFO:40:0.9 MINLEN:40’. The Nanopore sequencing reads (coverage >700x) were base-called with Guppy v5.0.0 [25] via MinKNOW v21.11.8 and corrected using Necat v0.0.1 [31] with ‘PREP_OUTPUT_COVERAGE=100 CNS_OUTPUT_COVERAGE=50’ options then assembled using Flye v2.6 [26] with default parameters. The initial assembly was corrected using polished Nanopore sequencing reads by racon [32] twice with ‘-m 8 -x -6 -g -8 -w 500’ options and the Illumina sequencing reads by Pilon v1.24 [33] three times with default parameters. The assembled contig was closed as validated using Bandage v0.8.1 [29].

### Taxon selection and phylogenomics

To expand the taxon selection for the CHAB-I-5 clade, we downloaded Rhodobacterales genomes from the NCBI and IMG databases (October, 2022), as well as large-scale metagenomic analyses including TARA Ocean [34, 35], BioGoShip [36], and OceanDNA [37]. First, a total of 259 genomes closely related to CHAB-I-5 (and a sister clade, represented by genomes like AG-337-I11 [38]) were selected based on having ANI values below 80% to other Roseobacters outside of these two groups. To categorically define the CHAB-I-5 cluster separate from the AG-337-I11 outgroup clade, 120 conserved bacterial single-copy genes were extracted and aligned using GTDB-tk v1.7.0 [39], and a phylogenetic tree was then constructed using IQ-TREE v2.2.0 [23] with parameters “-m LG+I+G -B 1000” (**Fig. S2**). We then dereplicated redundant CHAB-I-5 genomes using dRep v3.2.0 [40] with option ‘-pa 0.99 -ps 0.99’, which sets average nucleotide identity at 99%. Genomes with higher estimated quality, which was defined as completeness minus five times the amount of contamination [41], were selected as representatives for the recruitment analysis. We also excluded one genome, OceanDNA_b28631, because of its occurrence on a long branch in the phylogenomic tree and very low ANI (see below) to the remaining CHAB-I-5 genomes, which made its membership in this cluster questionable (**Fig. S3**). The resulting set included 52 representative CHAB-I-5 genomes, which were used to build the final phylogenomic tree using the same approach described above (**Fig. 1**). This phylogenomic tree was rooted using mad v2.2 based on minimal ancestor deviation approach [42]. This approach considers each branch as a possible root position, evaluates the ancestor-descendant relationships of all possible ancestral nodes in the tree, and chooses the branch with the minimal relative deviation as the root node [42].

**Figure 1.**
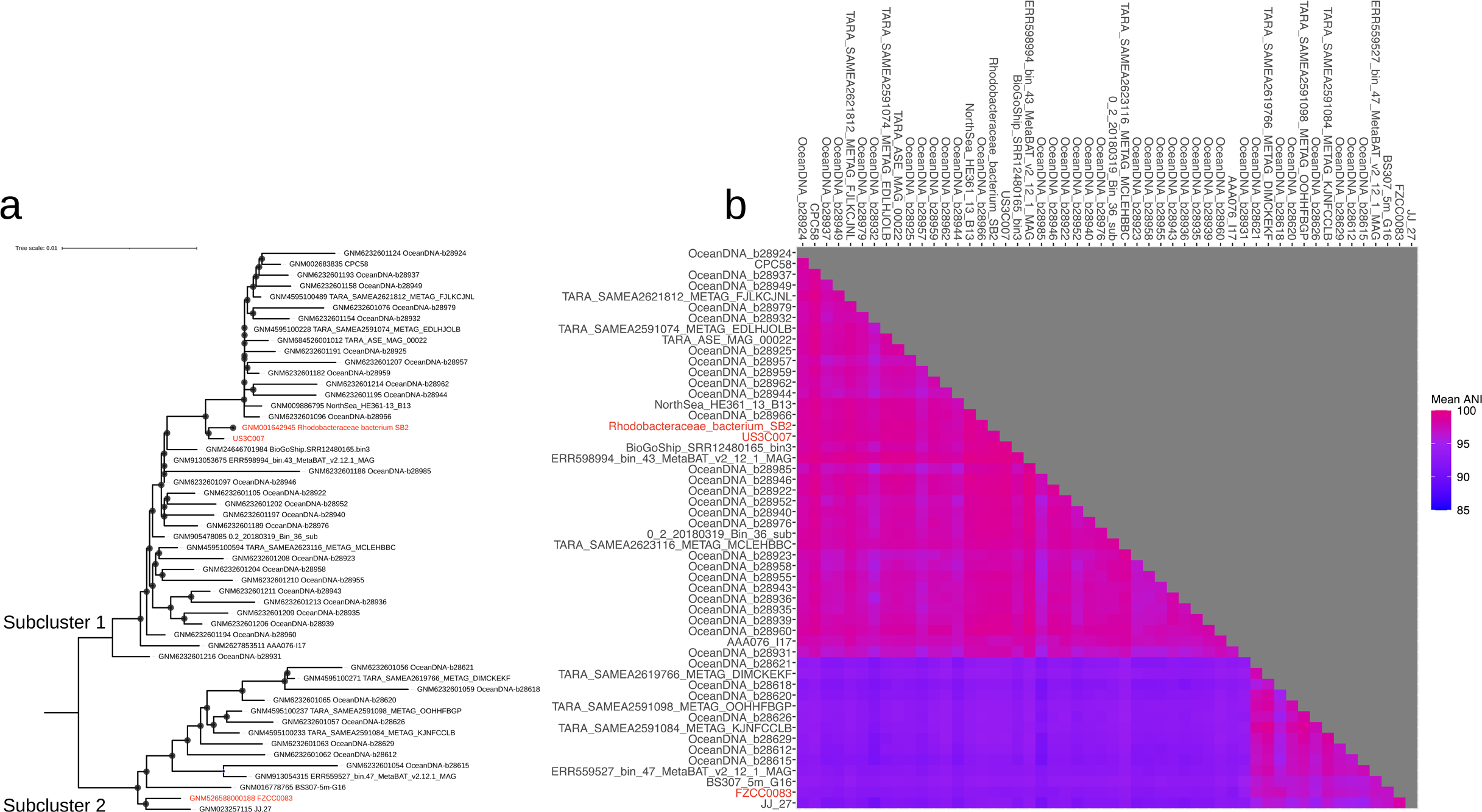
Phylogenomics and average nucleotide identity (ANI) of CHAB-I-5. **A)** Phylogenomic tree of 52 CHAB-I-5 genomes rooted with minimal ancestor deviation. CHAB-I-5 isolates are highlighted in red and Subclusters are labeled. Scale bar indicates changes per position. Filled circles indicate nodes with bootstrap values ≥ 95%. **B)** Pairwise ANI of the CHAB-I-5 Subclusters, colorized according to the key. Squares denoting 100% identity of each genome to itself are not colored.

### Comparative genomics

We compared the pairwise average nucleotide identity (ANI) with fastANI v1.33 [43] and visualized it in R. We used CheckM v1.1.3 [41] to evaluate all genomes and the specific ssu_finder function to identify the bacterial 16S rRNA genes. NCBI BLASTn was used for pairwise 16S rRNA gene comparisons. We also analyzed the metabolic potential of the final 52 genomes using Anvio’ v7.1 [44] to generate predicted amino acid sequences from genome sequences and GhostKOALA [45] for annotation of the amino acid sequences with the KEGG orthology database [46]. The resulting annotations and the original amino acid sequences were used with KEGG-Decoder and KEGG-Expander v.1.3 [47] to catalog the metabolic pathways present (**Fig. 3**). These metabolic annotations were further validated by searching against reference Roseobacter genomes (including *Ruegeria pomeroyi* DSS-3, *Dinoroseobacter shibae* DFL12, and *Planktomarina temperata* RCA23) using Orthofinder v2.2.1 [48]. These KEGG comparisons for all genomes are included in **Table S1**.

### Metagenomic read recruitment

Using 1,425 metagenomic samples from Yaquina bay [49], Sapelo Island [50], San Pedro Channel [51, 52], Baltic Sea [53, 54], Chesapeake Bay [55, 56], Columbia River [57], Black Sea [58], Gulf of Mexico [59], Pearl River [60], San Francisco Bay [61], and the North Pacific Subtropical Gyre [62] along with globally distributed metagenomic datasets [63–67], we recruited reads to the CHAB-I-5 genomes using competitive read recruitment via RRAP (91) as previously reported (20). Briefly, RRAP uses the latest versions of Bowtie2 [68] and SAMtools [69] to perform a competitive read recruitment from metagenomic samples to genomes, sort and index mapped reads, and normalize the data into RPKM values (Reads Per Kilobase (of genome) per Million (of recruited read base pairs)). We then analyzed the output in R. The OceanDNA_b28631 was included in the recruitment with the other 52 genomes, but excluded from visualization since we excluded it from our comparative genomics. The RPKM values for all the genomes are in **Table S1**.

### Microscopy

We initiated sample preparation by growing the US3C007 culture to a density of up to 10^6^ cells/ml, ensuring they were in the exponential growth phase. Subsequently, we fixed the cells in 2.5% (final concentration) glutaraldehyde. To harvest the fixed cells, we passed 10 mL of the bacterial suspension through a 0.2 µm Isopore polycarbonate filter (MilliporeSigma) coated with poly-L-lysine, facilitating cell adhesion to the filter membrane. Poly-L-lysine coating was achieved by immersing the membrane filter in a solution of Sigma P92155 at a concentration of 0.1 mg/mL. For cell membrane staining, we utilized a solution comprising 0.1 M HEPES buffered 0.05% Ruthenium Red (RuRed) with 10% sucrose. 10 mL of the RuRed solution was slowly passed through the filter, allowing for a 10-minute incubation period to ensure thorough staining. Then we performed a staining-fixing step using a solution containing 0.1 M HEPES buffered 0.05% RuRed, 0.8% Osmium tetroxide, and 10% sucrose. Similar to the previous step, 10 mL of the solution was slowly passed through the filter, with cells incubated for a minimum of 25 minutes. Following the staining-fixing process, we washed the membrane filter sequentially with two solutions: 10 mL of 0.1 M HEPES with 10% sucrose, followed by 10 mL of deionized water. Each washing step was carried out over a 10-minute duration. Subsequently, the samples underwent sequential dehydration in 50%, 70%, 95%, and 100% ethanol. The filter membrane was transferred to corresponding ethanol solutions in small containers and subjected to microwave treatment for one minute in a laboratory microwave oven, maintaining a temperature below 40°C. Finally, the samples were preserved in 100% ethanol on the filter and stored at room temperature for further analysis. The filters were sputtercoated for 45s with a Cressington 108 and imaged with the JSM-7001F-LV scanning electron microscope at the University of Southern California Core Center of Excellence in NanoImaging (https://cni.usc.edu). Resulting images were analyzed as described previously [70].

### Cell size analyses

Here we used a method adapted from previous studies [70]. Briefly, we segmented the cell image into two half spheres and a curved cylinder, mimicking a capsule geometry. The cell volume was then calculated as the sum of the volumes of the curved cylinder and the two half-spheres. While an ideal capsule assumes uniform radii for the half-spheres and the curved cylinder, variations in radii across different cell sections were addressed by measuring radii at multiple points and calculating geometric parameters (surface areas, volumes, lengths) based on each radius. Mean and median values of these parameters were used for visualization in our final violin plots (**Fig. 5A**).

We use Concepts for iPad v6.13 to measure to scale the image and to manually segment the cell area, measured in pixel squared (S). Additionally, the ruler feature in the application was employed to measure the radii of the cell by drawing circles covering widths at various sections, with each circle’s radius recorded as ‘r’. Mathematical equations for calculating surface areas (SA), volumes (V), lengths (l), and height (h) based on S and r are detailed in **Fig. S9**. Our analysis encompassed the geometries of 24 cells, as depicted in **Figs. S9** and **S10**, where we also showcase the segmentations and circles drawn for measurements.

### Growth experiments

The carbon substrate experiment for US3C007 was completed first using modified versions of the isolation medium, AMS1 (**Table S1**), to adjust the carbon concentrations while keeping all other components of the isolation medium the same. We tested the following concentrations of carbon serving as the presumptive electron donor and carbon source: 0, 9.37, 18.8, 37.5, 75, 150, 300, and 600 µM, as a 1:5:5 molar ratio mixture of methionine, glycine, and pyruvate (**Table S1**). For better cell yields, we then further modified the media compositions in AMS1 and created a new recipe, CCM (**Table S1**). In brief, the CCM media has no sulfate, completely relies on methionine for reduced sulfur, and has a 20X concentrated vitamin mix compared to AMS1. In addition, we substituted asparagine instead of glycine (which was added to aid in culturing SAR11 [17]) based on the genomic prediction of asparagine auxotrophy. We calculated the salinity of our media based on the chlorinity (salinity (ppt) = 1.80655 x Cl (ppt) [71]) of the “base salts”. Although a small amount of chloride also comes from our nutrients (e.g. ammonium chloride as the nitrogen source, manganese chloride and nickel chloride in the trace metals), the concentrations were negligible and we therefore did not incorporate those into the calculation of chlorinity. To test the salinity range of US3C0007, we made two batches of CCM at 0 and 50 ppt salinity and mixed them in different proportions (**Table S1**) to obtain the following salinities (ppt): 0, 5, 10, 15, 20, 25, 30, 35, 40, 45, and 49. The salinity experiment was conducted at 18.5°C (room temperature at that time). The temperature experiment was conducted using the CCM 30 ppt salinity medium at the following temperatures (°C): 4, 12, 16, 18.5, 21, 25, 28.5, 30, and 35.

Our methods of cell enumeration also evolved. For the carbon experiments, we counted cells with flow cytometry after staining with 1x Sybr green (Lonza 50513) for 30 minutes in the dark. The carbon experiments were enumerated on the Guava Easy Cyte 5HT flow cytometer (Millipore, Massachusetts, USA) as described above, except for the third transfer (fourth growth cycle) of the carbon concentration experiment, which was enumerated with an Accuri C6 Plus (Becton Dickinson, New Jersey, USA). The cell signals on the flow cytometry were gated based on the scatter plot of forward scatter vs. green fluorescence area. For the salinity and temperature experiments, we stained the cells using a final concentration of 10X Sybr green in addition to 10X Tris-EDTA (Sigma-Aldrich T9285) and 0.25% glutaraldehyde (Sigma-Aldrich 354400). Tris-EDTA maintains the nucleotide staining reaction at pH 8. Glutaraldehyde helps fix the cells and permeabilize the membrane. Together with the more concentrated Sybr green, the additional pH buffer and fixative help improve the staining performance. For enumeration, we counted 30 μL–100 μL of sample/staining cocktail mixture using a medium flow rate (35μL/min, 15 μm core size), threshold (triggering channel) of green fluorescence (533/30 nm) intensity at 1000. The cell signals were gated based on the green (533/30 nm) vs. yellow (585/40 nm) fluorescence intensity scatter plot. Based on the emission spectrum of Sybr green-stained DNA, we gated the signals with a ratio around 10:3 for green vs. yellow fluorescence intensities. We excluded autofluorescence signals from debris or media components, which usually have significantly higher yellow or red fluorescence compared to Sybr green-stained cells.

Culturing experiments for FZCC0083 were completed with an AMS1-based medium supplemented with a modified mixed carbon source [72], (1× concentrations of carbon mixtures were composed of 0.001% [wt/vol] D-glucose, D-ribose, methionine, pyruvic acid, glycine, taurine, N-acetyl D-glucosamine, and 0.002% [vol/vol] ethanol). This medium was used at the following temperatures (°C): 4, 12, 16, 20, 24, 26, 30, 37. For the salinity experiment, we also used modified versions of the AMS1. We kept the concentration of all added nutrient stocks constant and changed salinity by diluting or increasing the salt stocks while keeping the ratio of components constant. The exception to this was for sodium bicarbonate, which we kept constant to maintain buffering capacity. We tested the following salinities (ppt): 1, 2.3, 4.5, 9, 18, 22.6, 25.8, 30.1, 36, 43 at 24°C in the dark. Cells were enumerated on a Guava EasyCyte 5HT flow cytometer as described above.

### Growth curve analysis

Growth rates were calculated using a method adapted from our previously published sparse-growth-curve [73]. First we applied a sliding window for every three time points and generated a linear regression of the time vs. log2 transformed cell densities using SciPy package (1.13.0). The slope of the linear regression gives us the instantaneous doubling rate. To fully capture the uncertainties and variation of the statistics, we assigned each of the estimated slopes, plus and minus the standard deviation, to the start, middle, and end of the sliding window. This gave us nine candidate instantaneous doubling rates from any three time point cell densities. Since the end of a sliding window would become the middle of the next sliding window, and the start of the next, etc., each unique time point contributes to multiple estimated growth rates. We took the median estimated growth rate for each unique time point. We used an automated method to identify the instantaneous growth rates belonging to exponential phase. We attempted to fit a sigmoid decay curve to the time vs. instantaneous doubling rate data with the expectation that the exponential phase would correspond to the period before the inflection point. If the curve fit failed, we took the top three instantaneous doubling rates with maximum absolute values. To demonstrate this growth rate calculation method, we have added an example iPython notebook at GitHub (https://github.com/thrash-lab/insta_growth).

### Spectrophotometry

We attempted to measure bacteriochlorophyll in strain US3C007 via spectrophotometry of *in vivo* (whole cells) and pigment extracts. We performed direct *in vivo* measurements of culture volumes ranging from 50mL to 1L and cell densities from mid 10^5^ to mid 10^6^ cells/mL. Some runs involved filtering the cells onto sterile 0.1 µm polyethersulfone (PES) Supor filters (PALL Corporation, Port Washington, NY, USA) or centrifugation of cells at 9,000 rpm for 30 minutes. We used sterile CCM2 media for blanks and references. We also performed a washed cell-suspension using 950mL of US3C007 culture filtered onto a sterile 0.1 µm PES Supor filter with a 100mL wash of carbonless artificial seawater media (YBC) [74] and resuspension in 2mL of 1x PBS. In this case, 1x PBS was used as the reference and blank. For both in vivo approaches, cells were placed in a quartz cuvette (Hellma GmbH & Co. KG, Müllheim, DEU) and analyzed on a SpectraMax M2 plate reader (Molecular Devices, San Jose, CA, USA). The settings used were “Absorbance”, 350 - 900nm and 700 - 840nm, to obtain a full and detailed spectra profile. We performed extract measurements by first filtering 500mL of US3C007 culture (5×10^5--^ cells mL^−1^; sterile 0.1 µm PES Supor). Following previous methodology [75], the PES filter with the cells was extracted in 2 mL of 100% EtOH using the following minor modifications: PES filters were incubated for ∼24 hours in sealed borosilicate glass tubes (VWR International LLC, Radnor, PA, USA). Following extraction, the 2mL solution was then centrifuged at 5,000 rpm for 5 minutes and 700µL of supernatant was placed in a quartz cuvette for analysis with the plate reader as described above. 100% EtOH was used as the blank and reference. All samples were kept in the dark or wrapped in foil to prevent BChla degradation.

## Results

### Isolation, Identification, and Genome Sequencing

US3C007 originated from a cultivation experiment inoculated with surface water collected from the San Pedro Ocean Time series (SPOT) monthly cruise on 16 September 2020. Its top 16S rRNA gene BLAST hit was 100% identity to Roseobacter sp. SB2, accession KX467571.1 [7]. The 16S rRNA gene phylogeny at the time of isolation indicated US3C007 was the nearest phylogenetic neighbor to SB2 and the original clone library sequence of CHAB-I-5 [20] (**Fig. S1**). Strain FZCC0083 was isolated from the coastal waters of the East China Sea as previously described [10]. Hybrid long and short read genome sequencing resulted in single circularized contigs for both strains. Statistics for both genomes are reported in **Table 1** in comparison to the previously isolated strain SB2 [7]. All three strains have very similar sizes, GC content, and coding densities.

**Table 1.** CheckM genome statistics for the current and previously isolated strains.

| Genome | Length (bp) | Scaffolds (circular) | N50 (bp) | GC % | Coding genes | Coding density | # rRNA gene operons | Reference |
| --- | --- | --- | --- | --- | --- | --- | --- | --- |
| US3C007 | 3,622,411 | 1 (y) |  | 50.7 | 3,513 | 0.88 | 2 | This study |
| FZCC0083 | 3,646,439 | 1 (y) |  | 50.5 | 3,564 | 0.88 | 2 | This study |
| SB2 | 3,636,317 | 38 (n) | 323,631 | 50.5 | 3,527 | 0.89 | 1 | [7] |

Phylogenomic analysis of 52 CHAB-I-5 genomes resulted in two subgroups, with US3C007 and SB2 on one branch and FZCC0083 on the other (**Fig. 1A**). We refer to these two branches as Subcluster 1 and Subcluster 2, respectively. Subcluster 1 had a minimum within-cluster average nucleotide identity (ANI) of 95.2%, whereas Subluster 2 had a minimum within-cluster ANI of 94.5%. Between-cluster ANI percentages decreased below the species boundary, with a minimum of 90.6%, matching the phylogenomic branching pattern (**Fig. 1**). US3C007 and FZCC0083 both had two copies of the 16S rRNA gene. The two from US3C007 had 100% identity with SB2 (accession KX467571.1) and the two from FZCC0083 had 99.91% identity with SB2. The SB2 genome had only one copy of the rRNA gene operon located on a short contig, suggesting the other copy might not have assembled successfully. Comparisons of bulk genome characteristics across all 52 genomes showed a strong conservation of GC content (51.3 ± 0.4 %) and predicted coding density (89.3 ± 0.9 %) within CHAB-I-5 (**Table S1**).

### Biogeography

We mapped metagenomic reads to the CHAB-I-5 genomes from over fourteen hundred samples spanning a wide biogeography, including large ranges of salinity and temperature, to quantify CHAB-I-5 distribution. CHAB-I-5 was cosmopolitan, recruiting reads from around the globe. US3C007 was one of the top three most abundant representative genomes, including FZCC0083 and the ERR559527_bin_47_MetaBAT_v2_12_1_MAG as the first and second most abundant, with AA076_I17 and SB2 rounding out the top five (**Fig. 2A**). We observed recruitment across all latitudes and saw no specific relationship between genomes and latitude separate from that conferred by the locational bias of the samples themselves (**Figs. S4, S7**). Comparison of read recruitment with salinity demonstrated that all members of CHAB-I-5 prefer marine habitats, though some genomes do recruit limited numbers of reads from samples with a salinity as low as 8 (**Figs. S5, S7**). We also observed a tendency for genomes to recruit more reads from samples between temperatures of 11-20°C, even though most samples were from warmer locations (**Figs. S6, S7**). When abundance was summed by phylogenetic subcluster, the median recruited reads were smaller for Subcluster 1 than that of Subcluster 2 (**Fig. 2B**). However, Subcluster 1, containing isolates US3C007 and SB2, had a higher recruitment than Subcluster 2, the FZCC0083 type, at sites such as the Western United States coast, the Western South African coast, the North Sea, and the English Channel (**Fig. 2B**). Cluster 2 had higher recruitment at locations such as the Mediterranean, Pearl River, and much of the North Atlantic Gyre.

**Figure 2.**
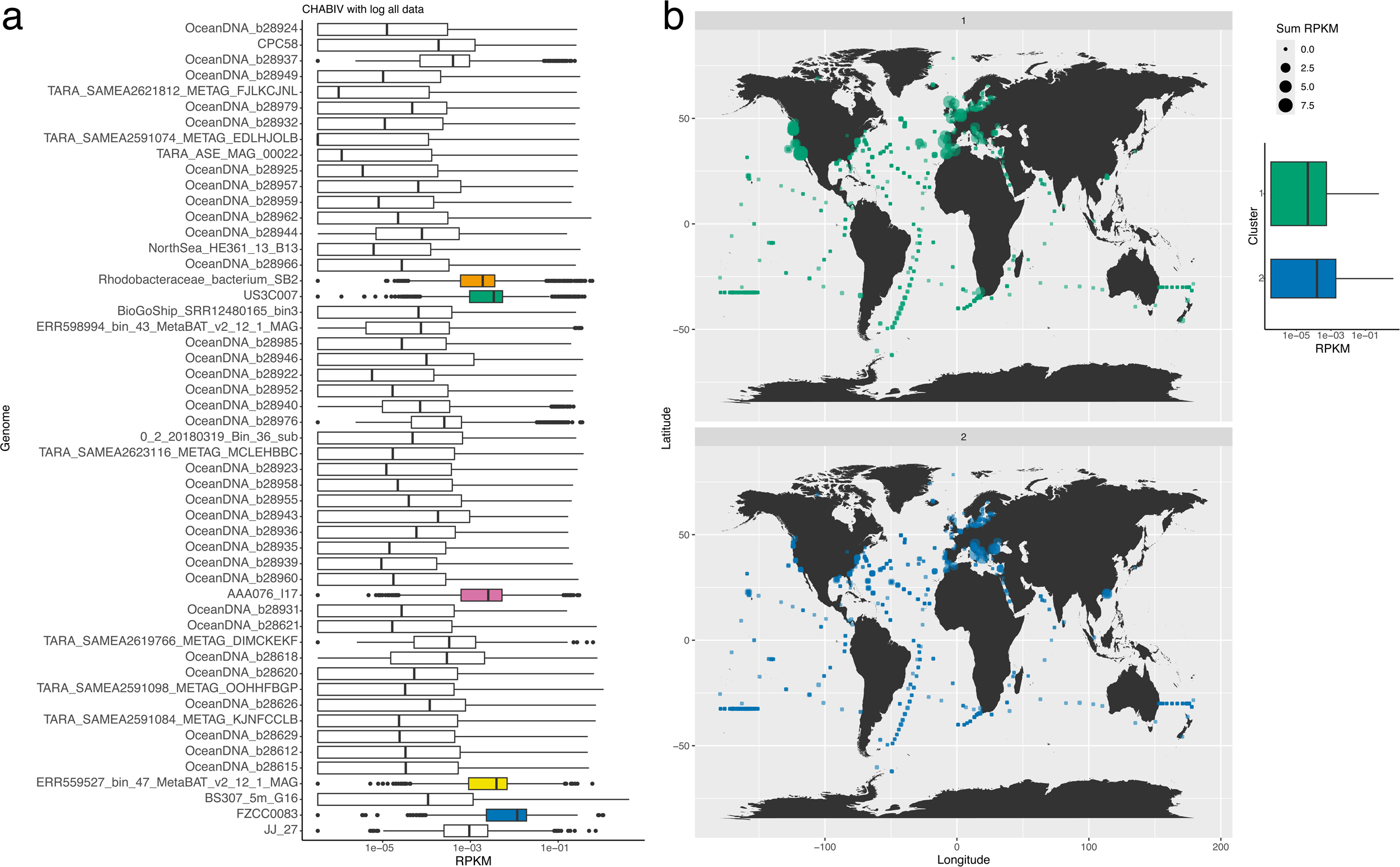
Biogeography and prevalence of CHAB-I-5 representatives. **A)** Boxplots of all RPKM values for each genome in the analysis. Black lines within the boxes indicate median RPKM values. The top five recruiting genomes are colored. **B)** Summed RPKM values for all genomes in each Subcluster, plotted according to sample location. RPKM values are depicted by circle size according to the key. Boxplot indicates the range of values for all genomes in each Subcluster.

### Genomic content

We compared CHAB-I-5 genomes to determine the conservation of metabolic potential and whether the two subclusters could be distinguished genomically (**Fig. 3**, **Table S1**). Corroborating previous reports [7, 12], these organisms were predicted to be capable of aerobic chemoorganoheterotrophic metabolism with the potential for anoxygenic phototrophy. All of the 52 non-redundant genomes had the potential for glycolysis via the Entner-Doudoroff pathway and the TCA cycle. All genomes contained nearly or fully complete electron transport pathways consisting of NADH-quinone oxidoreductases, F-type ATPases, cytochrome c oxidases, and ubiquinol-cytochrome c reductases. One genome, CPC58, contained a predicted *cbb_3_*-type cytochrome c oxidase. Most genomes had genes for polyhydroxyalkanoate (PHA) synthesis, a partial formaldehyde assimilation pathway, and a di/tri methylamine dehydrogenase. Most genomes had the potential to convert ethanol to acetate and acetaldehyde, and genes for anaplerotic C-fixation. Most genomes also contained a complete anoxygenic type-II reaction center. We found predicted genes for synthesis of bacteriochlorophyll a and/or b (*bchXYZ*, *bchC*, *bchF*, *chlG*, *chlP*-situated near the *puf* gene operon in US3C007) conserved across the CHAB-I-5 group, but found no annotated homologs for synthesis of bacteriochlorophyll d, c, or e. Full or partial pathways for flagella were also conserved.

**Figure 3.**
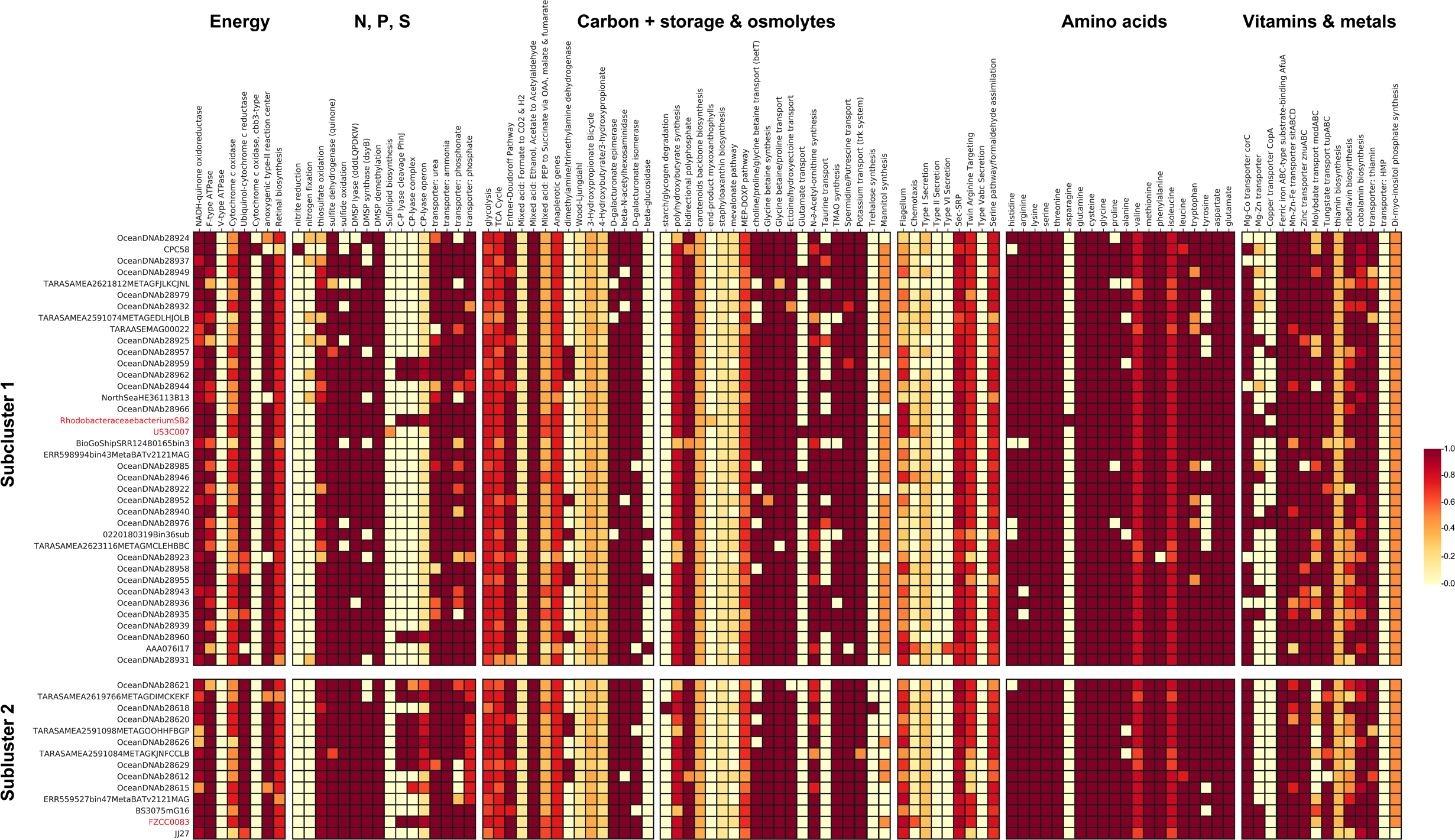
Predicted metabolism of CHAB-I-5. Subclusters are organized top to bottom to match the phylogeny of **Fig. 2**. Colors inside boxes correspond to pathway completion percentage according to the key. Genomes from isolates are noted in red.

Genes for metabolism of nitrogen, sulfur, phosphorous, trace metals, and vitamins were largely similar between subclusters. We found transporters for urea, ammonium, and phosphate were conserved, and most genomes contained a phosphonate transporter (**Fig. 3**). All 52 non-redundant genomes had the *napA* nitrate reductase, and CPC58 was the sole genome to encode a *nirK* nitrite reductase. All genomes except for CPC58 contained nearly or full pathways for thiosulfate oxidation via the *sox* gene cluster, and a gene encoding a sulfite dehydrogenase quinone was conserved. All non-redundant genomes in Subcluster 1 and most in Subcluster 2 had genes for sulfide oxidation. Most genomes had a DMSP lyase, all had genes for DMSP demethylation and most genomes also encoded a DMSP synthase. Predicted urease genes were prevalent throughout both Subclusters. Several genomes also encoded the C-P lyase complex, operon, and cleavage potential, although the latter was more common in Subcluster 2, and the US3C007 genome did not encode for the C-P lyase. Most genomes had a predicted Mg-Co transporter, Mg-Zn transport potential, and some genomes in Subcluster 1, including US3C007, had a *copA* copper transporter. Most genomes had ferric iron, Mn-Zn-Fe, zinc, and tungstate transporters. Most genomes in Subcluster 1 contained partial or nearly complete pathways for molybdate transport whereas only one genome in Subcluster 2 contained at least a half pathway. No genome contained the full pathway for thiamin biosynthesis, though a partial pathway was common. Most genomes contained either a full or partial pathway for riboflavin and cobalamin biosynthesis and thiamin transport.

Amino acid metabolism was also very similar between subclusters. Prototrophy for lysine, serine, threonine, glutamine, histidine, arginine, cysteine, glycine, valine, methionine, isoleucine, tryptophan, aspartate, and glutamate was largely conserved, whereas asparagine auxotrophy was widespread (**Fig. 3**). Glycine betaine synthesis, glycine betaine/proline transport, and ectoine/hydroxyectoine transport were also conserved. Most genomes in Subcluster 1 could transport taurine, with no genomes from Subcluster 2 containing this pathway, including the complete genome of FZCC0083.

Thus, the genome content across both subclusters was remarkably similar. The notable differences between the subclusters were the presence of the taurine and copper transporters as well as *pcaGH* dioxygenase genes exclusively within Subcluster 1, a greater prevalence of C-P lyase genes and low affinity phosphate transporter in Subcluster 2. Therefore, the Subclusters within CHAB-I-5 may exhibit some niche differentiation based on dissolved organic nitrogen and phosphorus utilization.

### Physiology and morphology

US3C007 grew consistently between 16 - 25°C, but not at temperatures of 12°C or below, or at 28.5°C or above (**Fig. 4A**). Additionally, US3C007 grew at salinities of 15-49 ppt, but not at 10 ppt or below (**Fig. 4B**). The maximum observed growth rate was 1.55 +/− 0.05 divisions day^−1^ at 18.5°C and 30 ppt (**Fig. 4B; Table S1**). We tested US3C007’s growth across a range of carbon concentrations to determine the carbon concentration to which it was best adapted. The primary carbon sources in the carbon mix were methionine, glycine, and pyruvate at a 1:5:5 molar ratio. We tested eight concentrations up to 600 µM carbon, with 300 µM carbon being the concentration in AMS1 medium (resulting from 10 µM methionine, 50 µM glycine, and 50 µM pyruvate) (**Table S1**). We observed no net change in growth rate with increasing carbon concentration, but an increase in yield (**Fig. 4C,D**). The consistent growth rate at low carbon concentrations indicates that this strain is particularly well adapted to low carbon environments. However, even after three transfers (four total growth cycles from lag to late log or stationary phase), we observed continued growth in the negative control, (**Fig. 4C**), albeit to lower cell densities than those cultures receiving carbon additions (**Fig. 4D**). We hypothesize this growth resulted from the strain having genes for PHA storage (**Fig. 3**), but this remains to be tested. We attempted to measure bacteriochlorophyll in whole cells, but were unable to determine a definitive spectrophotometric peak. This could have been the result of inadequate biomass or growth conditions that did not lend themselves to bacteriochlorophyll production.

**Figure 4.**
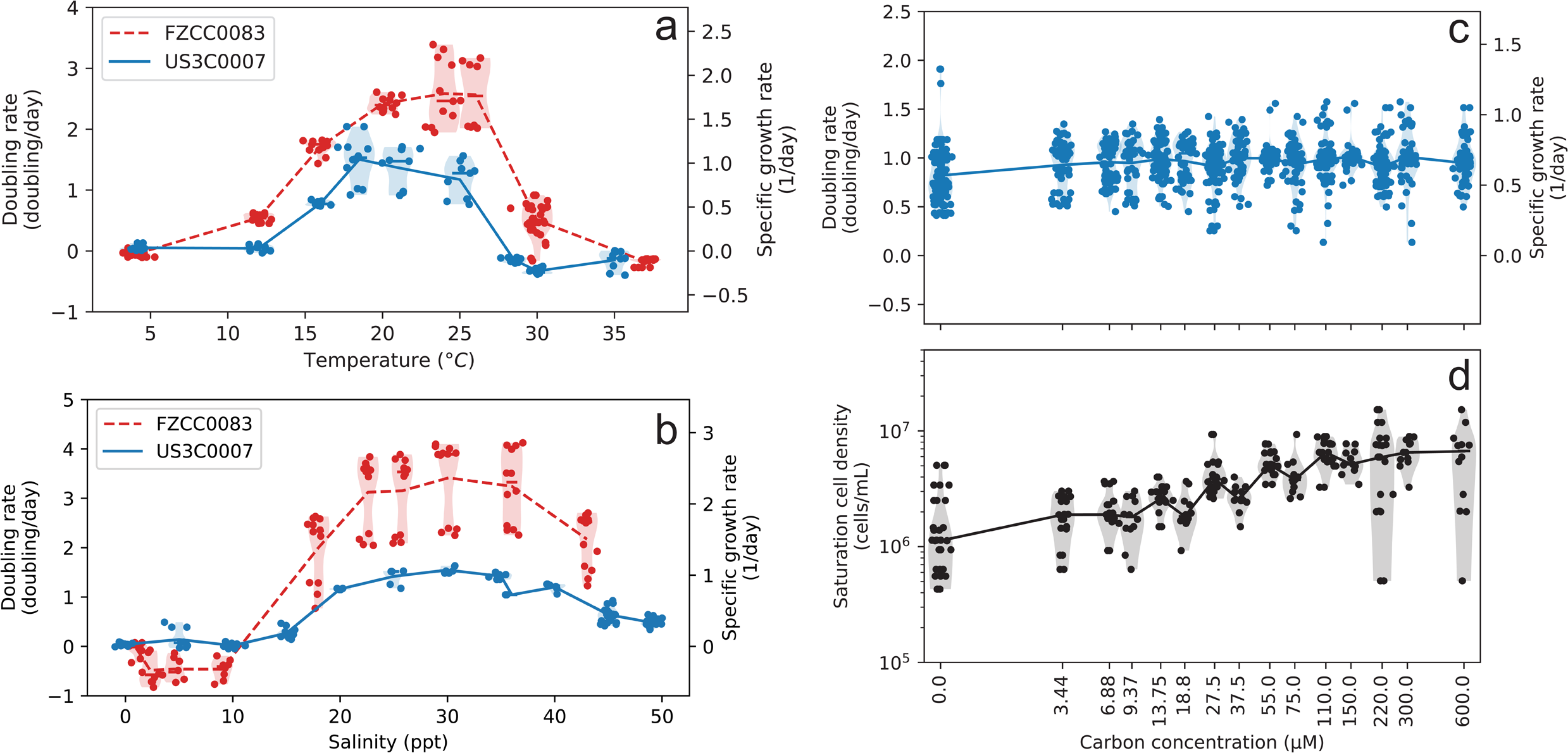
Growth rates for US3C007 and FZCC0083 across **A)** temperatures, **B)** salinities, and for US3C007 **C)** at differing low carbon concentrations. **D)** Growth yields for US3C007 under the same carbon experiments for C. Data in A-C comes from instantaneous growth rates throughout exponential phase. Data points are plotted along with medians (lines) and the distribution (violin plot shaded region).

FZCC0083 grew considerably faster than US3C007 in all conditions tested (**Fig. 4A,B; Table S1**). The maximum observed growth rate was 3.41 +/− 0.44 divisions day^−1^ at 24°C and 30.1 ppt (Fig. 4B)-more than twice the division rate of US3C007. FZCC0083 had a wider temperature growth envelope than US3C007, growing between 12 - 30°C with no growth at 4°C or 37°C. Its salinity tolerance was similar to that of US3C007, growing between 18-43 ppt, but not at 9 ppt or below. Thus, these two strains have notable differences in physiology which reflects their phylogenetic separation (**Fig. 1**).

US3C007 cells were small, having average cell lengths ∼1.65µm and radii ∼0.23 µm, yielding cell volumes ∼0.44 µm^3^ (**Fig. 5A**). We observed multiple morphologies within a single clonal culture (**Fig. 5B-E, Figs. S9, S10**). Single cells were usually bacillus-shaped, with some displaying more curved rod morphology or bulbous coccobacillus shapes (**Fig. 5B-E, Figs. S9-10**).

**Figure 5.**
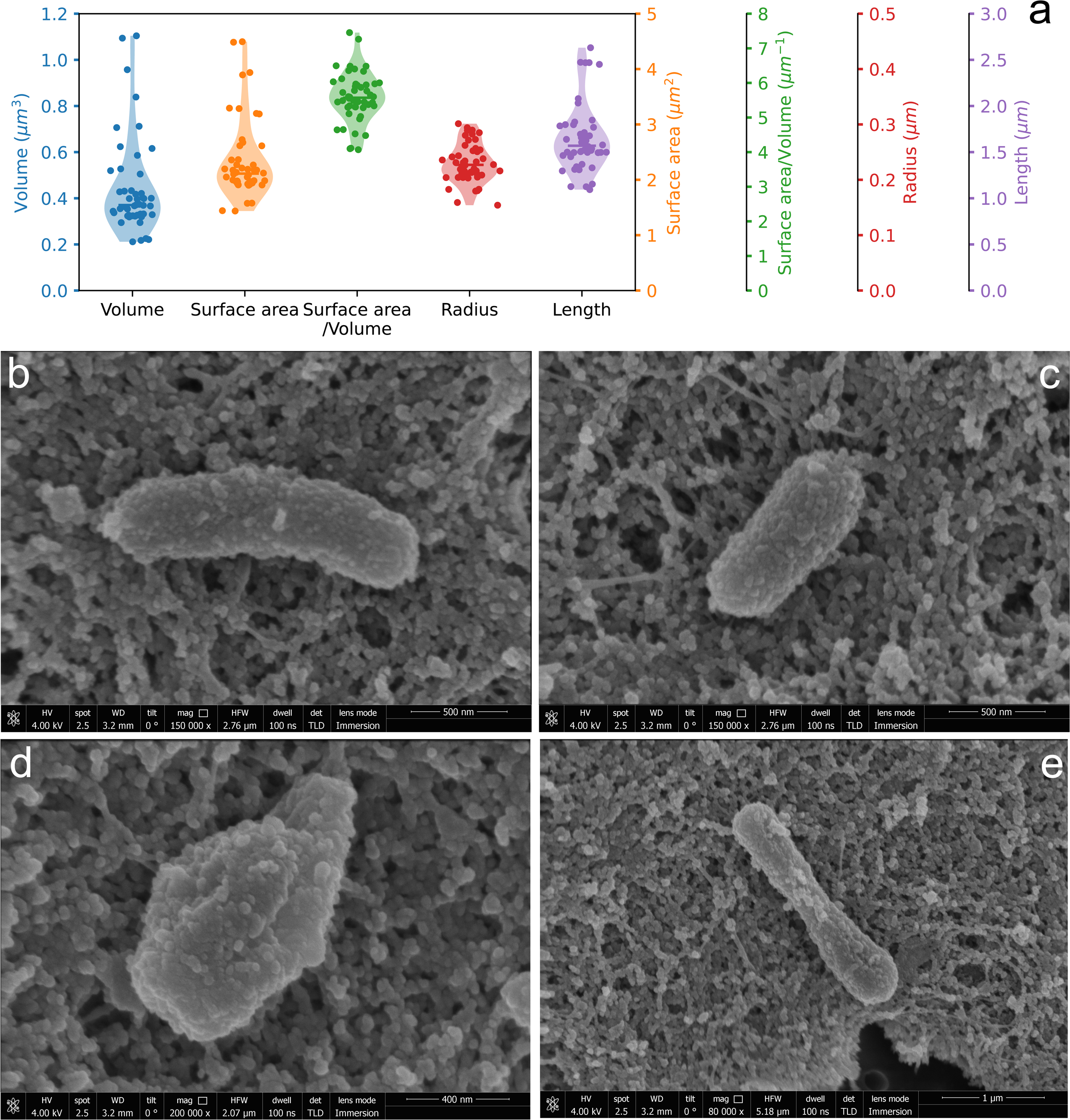
Cell size and shape of US3C007. **A)** Dimensions from analysis of 24 separate cells (see Figs. S9, S10) of different sizes and shapes. Medians are indicated with a bar and the violin plot shading shows the distribution of the data. **B-E)** Representative cells of different size/shape configurations seen in the culture. Scale bars (500 nm B,C; 400 nm D; 1 µm E) are indicated below each image.

## Discussion

This study is the most comprehensive analysis of the CHAB-I-5 subcluster within the larger “Roseobacter” group of *Rhodobacterales* to date. We have expanded the genomic and ecological characterization from four to 52 unique CHAB-I-5 genomes, including the first two circularized CHAB-I-5 genomes, and two new, publicly available CHAB-I-5 isolates, strains US3C007 and FZCC0083. Both strains are reliably propagated in artificial seawater media that are easily modified and we provided the first physiological and morphological characterization for members of the CHAB-I-5 group. Our expanded analysis also took advantage of recently generated, publicly available CHAB-I-5 genomes to understand intra-clade genomic diversity using phylogenomics and ANI. A prior study established two subclusters within CHAB-I-5 using environmental 16S rRNA gene sequence phylogeny [7], and we see the same division in our analysis. Both phylogenomics and average nucleotide identity support at least two subgroups within CHAB-I-5, denoted Subcluster 1 and Subcluster 2, that represent two species within a genus based on within- and between-subcluster ANI (**Fig. 1**). Isolate US3C007 belongs to Subcluster 1 and isolate FZCC0083 belongs to Subcluster 2.

Metabolic potential within CHAB-I-5 was highly conserved (**Fig. 3**). Nevertheless, we observed a few differences between Subclusters that may point to specific metabolic adaptations. The taurine ABC transporter *tauABC* was present in Subcluster 1 and not Subcluster 2 is (**Fig. 3**). Taurine is an important, multifunctional compound that serves as an osmoregulation tool and as a source for carbon, nitrogen, and sulfur for marine bacteria [76]. This differential ability to transport taurine may confer a growth advantage for Subcluster 1, but future research is needed to confirm how taurine is used, as all genomes encoded pathways for taurine catabolism. Another notable difference in metabolic content between the CHAB-I-5 subclusters was that of the C-P lyase genes. C-P lyases cleave carbon-phosphorous bonds and are used as a phosphate scavenging strategy that produces methane aerobically [77, 78]. Although all strains had typical *pstABCS* phosphate transporters, Subcluster 2 was enriched in C-P lyase genes, while only a few Subcluster 1 genomes had the pathway (**Fig. 3**). These results suggest Subcluster 2 interacts more consistently with the dissolved organic phosphorous pool and may contribute to methane production in global oceans. A subset of genomes in Subcluster 1 also contained the *copA* copper transporter exclusively, including US3C007, but distribution was spotty, suggesting a lack of conservation for the use of copper and/or that we didn’t observe the gene due to incomplete genomes.

We also extended the analysis of CHAB-I-5 distribution through read recruitment from over fourteen hundred metagenomic samples, including those in brackish and freshwater environments. Our results expand the known ecological distribution of CHAB-I-5 members, showing their presence in sample sites such as the North Atlantic gyre, South Pacific, Gulf of Mexico, Red Sea, and polar locations that were unavailable or had fewer sites surveyed in previous reports of CHAB-I-5 biogeography (**Fig. 2**) [7, 12]. Our work confirms and extends the view of CHAB-I-5 as a cosmopolitan member of the global oceans, and although the Subclusters were generally found in all the same locations, there were some samples where one Subcluster dominated (**Fig. 2B**). Subcluster 2 recruited more overall reads than Subcluster 1 (**Fig. 2B**), and the FZCC0083 and US3C007 genomes recruited the first and third most reads across all the samples (**Fig. 2A**). This suggests that these genomes are highly representative of CHAB-I-5 across the global oceans and make the strains excellent candidates for further study of the clade.

CHAB-I-5 can be abundant and active in polar latitudes [7, 12], however, our data did not show strong evidence of latitudinal preferences by genome (**Fig. S4**). Our initial physiological findings demonstrated restricted temperature range for both strains, representing each subcluster. US3C007 grew between 16-28.5°C and FZCC0083 grew between 12-30°C. These ranges are narrower than the observed range in metagenomic data for each genome, which both had substantial read recruitment in samples where the water temperatures were below 10°C (**Fig. S7**). This suggests the presence of (still uncultured) strains closely-related to US3C007 and FZCC0083 with greater psychrotolerance. In fact, many of the other SAG/MAG CHAB-I-5 genomes showed maximum read recruitment in samples below 15°C (**Fig. S6**), so it is likely that multiple strains of CHAB-I-5 are better cold-adapted than the two isolates.

On the other hand, the discrepancy between the lab and field measurements for US3C007 and FZCC0083 could describe the difference between realized and fundamental niches. Although the ideal fundamental niche space is sometimes envisioned as more extensive than the realized niche [79, 80], the reverse can also be true. For example, multiple ecotypes of *Prochlorococcus* had narrower temperature growth ranges in the laboratory than the ranges observed in nature via molecular data [81]. This is similar to the pattern we observed in both US3C007 and FZCC0083, where the realized niche appears larger than the fundamental niche with regards to temperature (**Figs. 4, S4**). For *Prochlorococcus*, the authors considered that a cultivation bias, stemming from continual maintenance of cultures in a restricted temperature range, could have led strains to evolve a different temperature optimum than the original population [81]. Similarly, continual culturing could also lead strains to evolve a more narrow temperature tolerance than would have been maintained by strains in the fluctuating natural environment. This would result in a contraction of the measured fundamental niche relative to the realized niche. However, another plausible explanation was that dispersal of *Prochlorococcus* resulted in cells being distributed to many locations outside their optimal temperature range [81]. Since our culture experiments are regularly restarted from cryopreserved samples, it was unlikely that our strains had evolved a more restricted temperature range since their isolation. Therefore, we consider our observations of a wider realized niche than fundamental niche to be consistent with the dispersal hypothesis as well.

Where salinity was concerned, our experiments suggested that CHAB-I-5 is an exclusively marine organism (**Fig. 4B**). Our read recruitment agreed - no genomes showed strong preferences for brackish or freshwater habitats (**Figs. S5, S7**). This contrasts with other abundant free-living microorganisms like SAR11 and Aegean-169 which both have subclades adapted to lower salinities [70, 82]. Other Roseobacter relatives have been isolated from brackish salinities as well [18, 19, 83, 84], and CHAB-I-5 members have been observed in equivalent abundances along the salinity gradient of the Chesapeake Bay [14]. While we found no clear evidence of fresh or brackish water specialists within CHAB-I-5, multiple genomes did recruit low numbers of reads from brackish waters with salinities as low as 8 (**Fig. S5**). Future work measuring activity of CHAB-I-5 across salinities could provide insight to whether the cells might be active in these lower salinity environments.

The US3C007 and FZCC0083 cultures have provided the first growth and morphological data for CHAB-I-5. These cultures span a wide range of growth rates, with the maximum for FZCC083 being over twice as fast as that of US3C007 (3.41 +/− 0.44 vs. 1.55 +/− 0.05 divisions day^−1^). These phenotypic differences likely reflect that these are different species, isolated from different oceanic regimes. Strain US3C007 was isolated from surface water collected at SPOT, a unique temperate semi-coastal location between Catalina Island and the coast of California overlying the San Pedro Basin at nearly 900m depth. Water circulation patterns in the Southern California Bight are complex [85, 86], but SPOT is inshore of the California Current system and average fall surface temperatures (warmest of the year) in the nearby Santa Monica Basin can reach 20.5°C [87]. Conversely, strain FZCC0083 was isolated from coastal waters off Pingtan Island, in the shallow Taiwan Strait very near the delineation of the East and South China Seas [10]. This location is in shallow water (< 30m) and over 8 degrees of latitude south of SPOT (∼900 km). Regional currents in this area branch from the Kuroshio Current system and fall average surface temperatures can reach 26°C [88]. Minimum temperatures in both areas are near 14°C. The optimization of FZCC0083 for growth at higher temperatures than US3C007, as well as the ability of FZCC0083 to grow at higher maximum temperatures (**Fig. 4A**), likely reflects the higher average temperatures in the Taiwan Strait compared to the Southern California Bight. Overall relative abundances of Subcluster 1 (US3C007-type) and Subcluster 2 (FZCC0083-type) with temperature were subtle, but showed general trends that match the isolate physiology: both trended downward with temperature, but Subcluster 1 had a slightly more negative correlation (**Fig. S11**). Thus, the growth physiology may signify larger habitat preferences for the Subclusters.

The considerable differences in growth rate between the two strains suggests more complex evolutionary diversification acting on multiple aspects of cell physiology. Nevertheless, the growth rates of these two strains span that of others in the larger PRC. Division rates for the model Roseobacter group organism, *Ruegeria pomeroyi* DSS-3^T^, which is not a PRC member, have been reported at up to 2.5 hour^−1^ [89]. The PRC includes many organisms with distinct genomic and lifestyle differences from better-studied copiotrophic Roseobacter group members like *R. pomeroyi* [7, 9, 90] and there are a few examples of cultured representatives from the PRC. Isolates from the DC5-80-3 (also called RCA) and CHUG groups that accompany CHAB-I-5 in the PRC have yielded some important growth insights [9, 91–93]. Strain LE17 had division rates of roughly 1 day^−1^ [92], and strain HKCCA1288 had division rates closer to 2 day^−1^ [9], although optimized medium has reduced this to just under 5 hours [89]. The type strain for the DC5-80-3 cluster, *Planktomarina temperata* RCA23^T^, as well as another close relative of US3C007, strain HIMB11, grew preferentially at mesophilic temperatures like US3C007, although rates were not reported [91, 94]. Given the close relationship between them, the variation in growth rates between US3C007 and FZCC0083 provide an excellent opportunity to investigate fundamental limits on growth rate. More strains from different locales will be important for exploring phenotypic heterogeneity within the group.

Our microscopic observation of strain US3C007 revealed significant pleomorphism in the culture (**Fig. 5B-E**). Pleomorphism and irregular morphology has been recorded in other Roseobacter group members, including HIMB11 [94]. Both HIMB11 and US3C007 have cells that are coccobacillus as well irregular rods [94] (**Fig. 5B-E**). The weighted average cell volume of 1,276 heterotrophic cells across 23 coastal ocean samples was 0.11 ± 0.17 µm^3^ [95], whereas average US3C007 volume was 0.44 ± 0.06 µm^3^ across a variety of morphologies (**Fig. 5A**). Thus, US3C007’s average cell volume is greater than the average heterotrophic bacterium, stemming in part from a relatively large radius for the cell length, compared to cells like that of SAR11 [70, 96, 97]. Future work to determine the extent of morphological variation and its drivers in natural populations of CHAB-I-5 will be important to understand the biology of these organisms more generally and for modeling the impact of carbon cycling by CHAB-I-5.

Overall, this work provides the most comprehensive genomic and ecological characterization of CHAB-I-5 and defines the first physiological data of the group. These recent advances in the availability of public CHAB-I-5 genomes and a new isolate that is representative of the CHAB-I-5 in global waters is a crucial component needed to characterize this abundant and highly active fraction of the microbial community. Future work is needed on US3C007 and the CHAB-I-5 cluster that could include comparative physiology between FZCC0083 and US3C007 to highlight whether a growth advantage might be conferred in the environment based on phosphorous, copper, or taurine availability and to quantify global estimates of CHAB-I-5’s contribution to biogeochemical cycling in the oceans.

### Description of *Thalassovivens*, gen. nov

*Thalassovivens* (Tha.las.so.vi’vens. Gr. fem. n. thalassa, the sea; L. pres. part. vivens, living, N.L. fem. n. Thalassovivens, an organism living in the sea, in reference to the marine habitat of these organisms)

Aerobic, with chemoorganoheterotrophic, chemolithotrophic, and anoxygenic phototrophic metabolisms. Encodes genes for glycolysis through the Entner-Doudoroff pathway and the TCA cycle. Genome sizes of ∼3.6 Mbp, with GC content ∼51% and a coding density ∼89%. Prototrophy predicted for lysine, serine, threonine, glutamine, histidine, arginine, cysteine, glycine, valine, methionine, isoleucine, tryptophan, aspartate, and glutamate, with asparagine auxotrophy. Glycine betaine synthesis, glycine betaine/proline transport, and ecotine/hydroxyectoine transport genomically conserved. Genes for the PII nitrogen regulatory system, *ntrXY*, *amtB*, and urease conserved. Most genomes also encode genes for aerobic vitamin B_12_ synthesis. Genes for synthesis of bacteriochlorophyll a and/or b conserved. Motility via flagella is predicted.

### Description of *Thalassovivens spotae*, sp. nov

*Thalassovivens spotae* (spo’tae. N.L. gen. n. spotae, in reference to the San Pedro Ocean Time series (SPOT), from which the strain was isolated).

In addition to the characteristics of the genus, it has the following features. Cells are coccobacillus shaped, pleomorphic, with average dimensions of 0.23 µm radius, 1.65 µm length, and 0.44 µm^3^ volume. Halotolerant, growing in salinities of 15-49 ppt, but not at 10 ppt or below. Mesophilic, growing between 16 −25°C, but not at temperatures of 12°C or below, or at 28.5°C or above. Has a maximum growth rate of 1.55 +/− 0.05 divisions day^−1^ at 20°C and salinity of 30 ppt.

The type strain, US3C007^T^, was isolated from surface water (2m) collected at the San Pedro Ocean Time series (33°33’ N, 118°24’ W). The genome sequence is circularized at 3,622,411 bp with 50.7% GC content. The genome is available on NCBI at BioProject number PRJNA1044073.

***Note to editors/reviewers: we sent strain US3C007 to both the DSMZ and ATCC culture collections in January 2024 and February 2024, respectively, and are awaiting confirmation of deposition. We would like to undergo review while the deposition process moves forward and we will update the accession numbers (ATCC XXXXX = DSMZ XXXXX) as part of our later revisions*.**

## Acknowledgments

We thank Dr. Aharon Oren for assistance with nomenclature and acknowledge the Center for Advanced Research Computing (CARC) at the University of Southern California for providing computing resources that have contributed to the research results reported within this publication (https://carc.usc.edu). This work was supported by a Simons Early Career Investigator in Marine Microbial Ecology and Evolution Award, and NSF Biological Oceanography Program OCE-1945279 and Emerging Frontiers Program EF-2125191 grants to J.C.T.

## Data Availability

The genome sequences and raw reads for strains US3C007 and FZCC0083 can be found at NCBI under BioProject numbers PRJNA1044073 and PRJNA1047292, respectively. Supplementary material, including scripts, tree files, and **Table S1**, can be found on FigShare 10.6084/m9.figshare.25898389.

## Supplemental Tables and Figures

**Table S1.** Excel spreadsheet containing genome statistics, computed ANI values, metabolic predictions, AMS1 medium recipe and modifications, growth rates for growth experiments, RPKM values from metagenomic recruitment, and microscopic size calculations. Table S1 is hosted at FigShare 10.6084/m9.figshare.25898389.

**Figure S1**. Phylogenetic tree of 16S rRNA gene sequences from the Alphaproteobacteria with US3C007 and other CHAB-I-5 representatives. Nodes outside of the CHAB-I-5 and Roseobacter HIMB11 clade have been collapsed to show US3C007’s inclusion with the CHAB-I-5 sequences. The CHAB-I-5 cluster is boxed in red and strain US3C007 is starred.

**Figure S2.** Phylogenomic tree of all CHAB-I-5 genomes prior to dereplication and those of the sister clade containing AG-337-I11 and others. Scale bar indicates changes per position. Filled circles indicate nodes with bootstrap values ≥ 95%.

**Figure S3.** Phylogenomic tree of dereplicated CHAB-I-5 genomes (excepting the dual copies of the SB2 genome), and associated ANI values. Dotted lines indicate the position of the OceanDNA_b28631 genome, which was removed due to the low ANI values and the long unsupported branch on the tree. Scale bar indicates changes per position. Filled circles indicate nodes with bootstrap values ≥ 95%.

**Figure S4.** Metagenomic recruitment (normalized as RPKM) to all genomes by latitude with non-linear regression lines featuring shading that represents the 95% confidence intervals. The histogram below the RPKM plots shows the sample distribution according to latitude.

**Figure S5.** Metagenomic recruitment (normalized as RPKM) to all genomes by salinity with non-linear regression lines featuring shading that represents the 95% confidence intervals. The histogram below the RPKM plots shows the sample distribution according to salinity.

**Figure S6.** Metagenomic recruitment (normalized as RPKM) to all genomes by temperature with non-linear regression lines featuring shading that represents the 95% confidence intervals. The histogram below the RPKM plots shows the sample distribution according to temperature.

**Figure S7.** Metagenomic recruitment (normalized as RPKM) for the top 5 recruiting genomes according to **A)** latitude, **B)** salinity, and **C)** temperature with non-linear regression lines featuring shading that represents the 95% confidence intervals. Histograms below the RPKM plots show the sample distribution according to the same x-axis variable. Note that while all metagenomic samples had latitude values, the metadata did not always include salinity or temperature, and thus the total number of points in B) and C) are different.

**Figure S8.** Growth curves of strains US3C007 and FZCC0083 for the temperature and salinity experiments. Y-axes are cell concentrations in cells/ml, x-axes are time. Conditions are written at the top of each plot.

**Figure S9.** Notes and marks for the analyses of cell morphologies. Using the pixel and scale features in Concepts for iPad v6.13, we measured the radii (R) and area of the cross section (S) of the cells. The formula of the lengths (l), volumes (V), and surface areas (SA) calculated based on r (we denoted r as the mean radii of each cell) and S are shown at the top of the figure. The detailed formula could also be found at **Table S1**.

**Figure S10.** Same as Figure S9, marks of measurements for the SEM images.

**Figure S11.** Relative abundance of Subclusters 1 (green) and 2 (blue) compared to temperature. Subcluster RPKMs were summed as in Figure 2B. R^2^ values for the linear regressions are plotted at the top. Shading around the linear regression indicates 95% confidence intervals.

