## Supplemental Figures for "New isolates refine the ecophysiology of the Roseobacter CHAB-I-5 lineage"

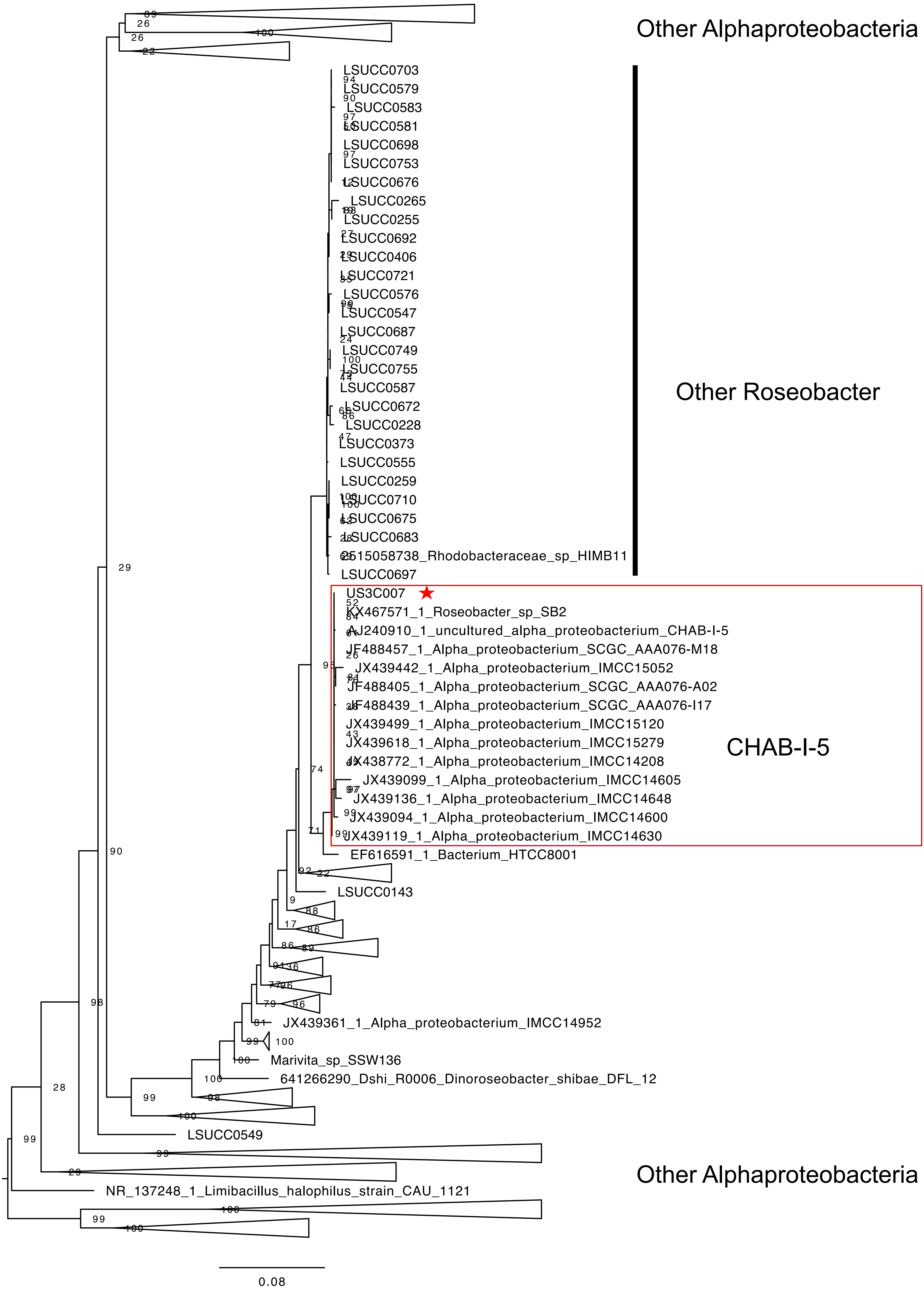

Other Alphaproteobacteria
LSUCC0703
LSUCC0579
LSUCC0583
LSUCC0581
LSUCC0698
LSUCC0753
LSUCC0676
LSUCC0265
LSUCC0255
LSUCC0692
LSUCC0406
LSUCC0721
LSUCC0576
LSUCC0547
LSUCC0687
LSUCC0749
LSUCC0755
LSUCC0587
LSUCC0672
LSUCC0228
LSUCC0373
LSUCC0555
LSUCC0259
LSUCC0710
LSUCC0675
LSUCC0683
2515058738_Rhodobacteraceae_sp_HIMB11
LSUCC0697
Other Roseobacter
US3C007
KX467571_1_Roseobacter_sp_SB2
AJ240910_1_uncultured_alpha_proteobacterium_CHAB-I-5
JF488457_1_Alpha_proteobacterium_SCGC_AAA076-M18
JX439442_1_Alpha_proteobacterium_IMCC15052
JF488405_1_Alpha_proteobacterium_SCGC_AAA076-A02
JF488439_1_Alpha_proteobacterium_SCGC_AAA076-I17
JX439499_1_Alpha_proteobacterium_IMCC15120
JX439618_1_Alpha_proteobacterium_IMCC15279
JX438772_1_Alpha_proteobacterium_IMCC14208
JX439099_1_Alpha_proteobacterium_IMCC14605
JX439136_1_Alpha_proteobacterium_IMCC14648
JX439094_1_Alpha_proteobacterium_IMCC14600
JX439119_1_Alpha_proteobacterium_IMCC14630
CHAB-I-5
EF616591_1_Bacterium_HTCC8001
LSUCC0143
JX439361_1_Alpha_proteobacterium_IMCC14952
Marivita_sp_SSW136
641266290_Dshi_R0006_Dinoroseobacter_shibae_DFL_12
LSUCC0549
Other Alphaproteobacteria
NR_137248_1_Limibacillus_halophilus_strain_CAU_1121
0.08

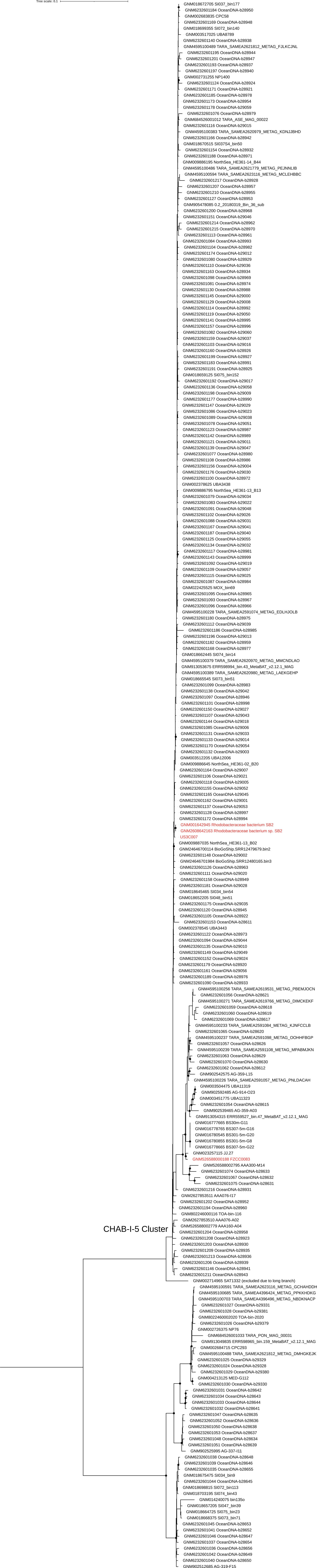

Tree scale: 0.1
GNM018672705 SI037_bin177
GNM6232601184 OceanDNA-b28950
GNM002683835 CPC58
GNM6232601169 OceanDNA-b28948
GNM018699355 SI072_bin140
GNM003517025 UBA8789
GNM6232601140 OceanDNA-b28938
GNM4595100489 TARA_SAMEA2621812_METAG_FJLKCJNL
GNM6232601195 OceanDNA-b28944
GNM6232601201 OceanDNA-b28947
GNM6232601193 OceanDNA-b28937
GNM6232601197 OceanDNA-b28940
GNM002731255 NP1400
GNM6232601124 OceanDNA-b28924
GNM6232601171 OceanDNA-b28921
GNM6232601185 OceanDNA-b28978
GNM6232601173 OceanDNA-b28954
GNM6232601178 OceanDNA-b29059
GNM6232601076 OceanDNA-b28979
GNM684526001012 TARA_ASE_MAG_00022
GNM6232601116 OceanDNA-b29015
GNM4595100383 TARA_SAMEA2620979_METAG_KDNJJBHD
GNM6232601166 OceanDNA-b28942
GNM018670515 SI037S4_bin50
GNM6232601154 OceanDNA-b28932
GNM6232601188 OceanDNA-b28971
GNM009886195 NorthSea_HE361-14_B44
GNM4595100486 TARA_SAMEA2621779_METAG_PEJNNLIB
GNM4595100594 TARA_SAMEA2623116_METAG_MCLEHBBC
GNM6232601217 OceanDNA-b28928
GNM6232601207 OceanDNA-b28957
GNM6232601210 OceanDNA-b28955
GNM6232601127 OceanDNA-b28953
GNM905478085 0.2_20180319_Bin_36_sub
GNM6232601200 OceanDNA-b28968
GNM6232601151 OceanDNA-b29046
GNM6232601214 OceanDNA-b28962
GNM6232601215 OceanDNA-b28970
GNM6232601113 OceanDNA-b28961
GNM6232601084 OceanDNA-b28993
GNM6232601104 OceanDNA-b28982
GNM6232601174 OceanDNA-b29012
GNM6232601080 OceanDNA-b28929
GNM6232601110 OceanDNA-b29036
GNM6232601163 OceanDNA-b28934
GNM6232601098 OceanDNA-b28969
GNM6232601081 OceanDNA-b28974
GNM6232601130 OceanDNA-b28988
GNM6232601145 OceanDNA-b29000
GNM6232601129 OceanDNA-b29008
GNM6232601114 OceanDNA-b28992
GNM6232601119 OceanDNA-b29050
GNM6232601141 OceanDNA-b28995
GNM6232601157 OceanDNA-b28996
GNM6232601082 OceanDNA-b29060
GNM6232601159 OceanDNA-b29037
GNM6232601103 OceanDNA-b29016
GNM6232601160 OceanDNA-b28926
GNM6232601199 OceanDNA-b28927
GNM6232601183 OceanDNA-b28991
GNM6232601191 OceanDNA-b28925
GNM018659125 SI075_bin152
GNM6232601192 OceanDNA-b29017
GNM6232601136 OceanDNA-b29058
GNM6232601198 OceanDNA-b29009
GNM6232601177 OceanDNA-b28990
GNM6232601147 OceanDNA-b29029
GNM6232601086 OceanDNA-b29023
GNM6232601089 OceanDNA-b29038
GNM6232601078 OceanDNA-b29051
GNM6232601123 OceanDNA-b28987
GNM6232601142 OceanDNA-b28989
GNM6232601121 OceanDNA-b29011
GNM6232601139 OceanDNA-b29047
GNM6232601077 OceanDNA-b28980
GNM6232601108 OceanDNA-b28986
GNM6232601156 OceanDNA-b29004
GNM6232601176 OceanDNA-b29030
GNM6232601100 OceanDNA-b28972
GNM002378625 UBA3438
GNM009886795 NorthSea_HE361-13_B13
GNM6232601079 OceanDNA-b29034
GNM6232601083 OceanDNA-b29022
GNM6232601091 OceanDNA-b29048
GNM6232601102 OceanDNA-b29026
GNM6232601088 OceanDNA-b29031
GNM6232601167 OceanDNA-b29041
GNM6232601187 OceanDNA-b29040
GNM6232601125 OceanDNA-b29055
GNM6232601134 OceanDNA-b29032
GNM6232601117 OceanDNA-b28981
GNM6232601143 OceanDNA-b28999
GNM6232601092 OceanDNA-b29019
GNM6232601109 OceanDNA-b29057
GNM6232601115 OceanDNA-b29025
GNM6232601087 OceanDNA-b28984
GNM022425525 MOX_bin69
GNM6232601095 OceanDNA-b28965
GNM6232601093 OceanDNA-b28967
GNM6232601096 OceanDNA-b28966
GNM4595100228 TARA_SAMEA2591074_METAG_EDLHJOLB
GNM6232601180 OceanDNA-b28975
GNM6232601112 OceanDNA-b29039
GNM6232601186 OceanDNA-b28985
GNM6232601196 OceanDNA-b29013
GNM6232601182 OceanDNA-b28959
GNM6232601168 OceanDNA-b28977
GNM018662445 SI074_bin14
GNM4595100379 TARA_SAMEA2620970_METAG_MMCNDLAO
GNM913053675 ERR598994_bin.43_MetaBAT_v2.12.1_MAG
GNM4595100389 TARA_SAMEA2620980_METAG_LAEKGEHP
GNM018665545 SI073_bin51
GNM6232601099 OceanDNA-b28983
GNM6232601138 OceanDNA-b29042
GNM6232601097 OceanDNA-b28946
GNM6232601101 OceanDNA-b28998
GNM6232601150 OceanDNA-b29027
GNM6232601107 OceanDNA-b29043
GNM6232601144 OceanDNA-b29018
GNM6232601085 OceanDNA-b29006
GNM6232601131 OceanDNA-b29033
GNM6232601133 OceanDNA-b29014
GNM6232601170 OceanDNA-b29054
GNM6232601132 OceanDNA-b29003
GNM003512205 UBA12006
GNM009886645 NorthSea_HE361-02_B20
GNM6232601164 OceanDNA-b29007
GNM6232601106 OceanDNA-b29021
GNM6232601118 OceanDNA-b29005
GNM6232601155 OceanDNA-b29052
GNM6232601165 OceanDNA-b29045
GNM6232601162 OceanDNA-b29001
GNM6232601137 OceanDNA-b29053
GNM6232601128 OceanDNA-b28997
GNM6232601172 OceanDNA-b28994
GNM001642945 Rhodobacteraceae bacterium SB2
GNM2608642163 Rhodobacteraceae bacterium sp. SB2
US3C007
GNM009887035 NorthSea_HE361-13_B02
GNM2646700114 BioGoShip.SRR12479679.bin2
GNM6232601148 OceanDNA-b29002
GNM2646701984 BioGoShip.SRR12480165.bin3
GNM6232601126 OceanDNA-b28963
GNM6232601111 OceanDNA-b29020
GNM6232601158 OceanDNA-b28949
GNM6232601181 OceanDNA-b29028
GNM018645465 SI034_bin54
GNM018652205 SI048_bin51
GNM6232601175 OceanDNA-b29035
GNM6232601120 OceanDNA-b28945
GNM6232601105 OceanDNA-b28922
GNM6232601153 OceanDNA-b28611
GNM002378545 UBA3443
GNM6232601122 OceanDNA-b28973
GNM6232601094 OceanDNA-b29044
GNM6232601135 OceanDNA-b29010
GNM6232601149 OceanDNA-b29049
GNM6232601152 OceanDNA-b29024
GNM6232601179 OceanDNA-b28920
GNM6232601161 OceanDNA-b29056
GNM6232601189 OceanDNA-b28976
GNM6232601090 OceanDNA-b28933
GNM4595100256 TARA_SAMEA2619531_METAG_PBEMJOCN
GNM6232601056 OceanDNA-b28621
GNM4595100271 TARA_SAMEA2619766_METAG_DIMCKEKF
GNM6232601059 OceanDNA-b28618
GNM6232601060 OceanDNA-b28619
GNM6232601069 OceanDNA-b28617
GNM4595100233 TARA_SAMEA2591084_METAG_KJNFCCLB
GNM6232601065 OceanDNA-b28620
GNM4595100237 TARA_SAMEA2591098_METAG_OOHHFBGP
GNM6232601057 OceanDNA-b28626
GNM4595100239 TARA_SAMEA2591108_METAG_MPABMJKN
GNM6232601063 OceanDNA-b28629
GNM6232601070 OceanDNA-b28630
GNM6232601062 OceanDNA-b28612
GNM902542575 AG-359-L15
GNM4595100226 TARA_SAMEA2591057_METAG_PNLDACAH
GNM003504475 UBA11319
GNM902592485 AG-914-O23
GNM003451775 UBA11323
GNM6232601054 OceanDNA-b28615
GNM902539465 AG-359-A03
GNM913054315 ERR559527_bin.47_MetaBAT_v2.12.1_MAG
GNM016777665 BS30m-G11
GNM016778765 BS307-5m-G16
GNM016780545 BS301-5m-G20
GNM016780855 BS301-5m-G8
GNM016778665 BS307-5m-G22
GNM023257115 JJ.27
GNM526588000188 FZCC0083
GNM526588002795 AAA300-M14
GNM6232601074 OceanDNA-b28633
GNM6232601067 OceanDNA-b28632
GNM6232601075 OceanDNA-b28631
GNM6232601216 OceanDNA-b28931
GNM2627853511 AAA076-I17
GNM6232601202 OceanDNA-b28952
GNM6232601194 OceanDNA-b28960
GNM802246000116 TOA-bin-116
GNM2627853510 AAA076-A02
GNM526588002779 AAA160-A04
GNM6232601204 OceanDNA-b28958
GNM6232601208 OceanDNA-b28923
GNM6232601203 OceanDNA-b28930
GNM6232601209 OceanDNA-b28935
GNM6232601213 OceanDNA-b28936
GNM6232601206 OceanDNA-b28939
GNM6232601146 OceanDNA-b28941
GNM6232601211 OceanDNA-b28943
CHAB-I-5 Cluster
GNM002714965 SAT1332 (excluded due to long branch)
GNM4595100591 TARA_SAMEA2623116_METAG_GCHAHDDH
GNM4595100685 TARA_SAMEA4396424_METAG_PPKKHDKG
GNM4595100703 TARA_SAMEA4396496_METAG_NBDKNACP
GNM6232601027 OceanDNA-b29331
GNM6232601028 OceanDNA-b29381
GNM802246002020 TOA-bin-2020
GNM6232601026 OceanDNA-b29379
GNM002726375 NP76
GNM684526001033 TARA_PON_MAG_00031
GNM913049835 ERR598965_bin.159_MetaBAT_v2.12.1_MAG
GNM002684715 CPC293
GNM4595100488 TARA_SAMEA2621812_METAG_DMHGKEJK
GNM6232601025 OceanDNA-b29329
GNM6232601024 OceanDNA-b29328
GNM6232601029 OceanDNA-b29380
GNM000213125 MED-G112
GNM6232601030 OceanDNA-b29330
GNM6232601031 OceanDNA-b28642
GNM6232601034 OceanDNA-b28643
GNM6232601033 OceanDNA-b28644
GNM6232601032 OceanDNA-b28641
GNM6232601047 OceanDNA-b28635
GNM6232601052 OceanDNA-b28636
GNM6232601050 OceanDNA-b28638
GNM6232601053 OceanDNA-b28637
GNM6232601048 OceanDNA-b28634
GNM6232601051 OceanDNA-b28639
GNM902525995 AG-337-I11
GNM6232601038 OceanDNA-b28648
GNM6232601039 OceanDNA-b28646
GNM6232601035 OceanDNA-b28655
GNM018675475 SI034_bin9
GNM6232601044 OceanDNA-b28645
GNM018698815 SI072_bin113
GNM018703195 SI074_bin43
GNM014240075 bin135o
GNM018657205 SI047_bin39
GNM018664725 SI075_bin23
GNM018668375 SI073_bin71
GNM6232601045 OceanDNA-b28653
GNM6232601041 OceanDNA-b28652
GNM6232601046 OceanDNA-b28647
GNM6232601037 OceanDNA-b28654
GNM6232601036 OceanDNA-b28656
GNM6232601042 OceanDNA-b28649
GNM6232601040 OceanDNA-b28650
GNM902512685 AG-319-F15

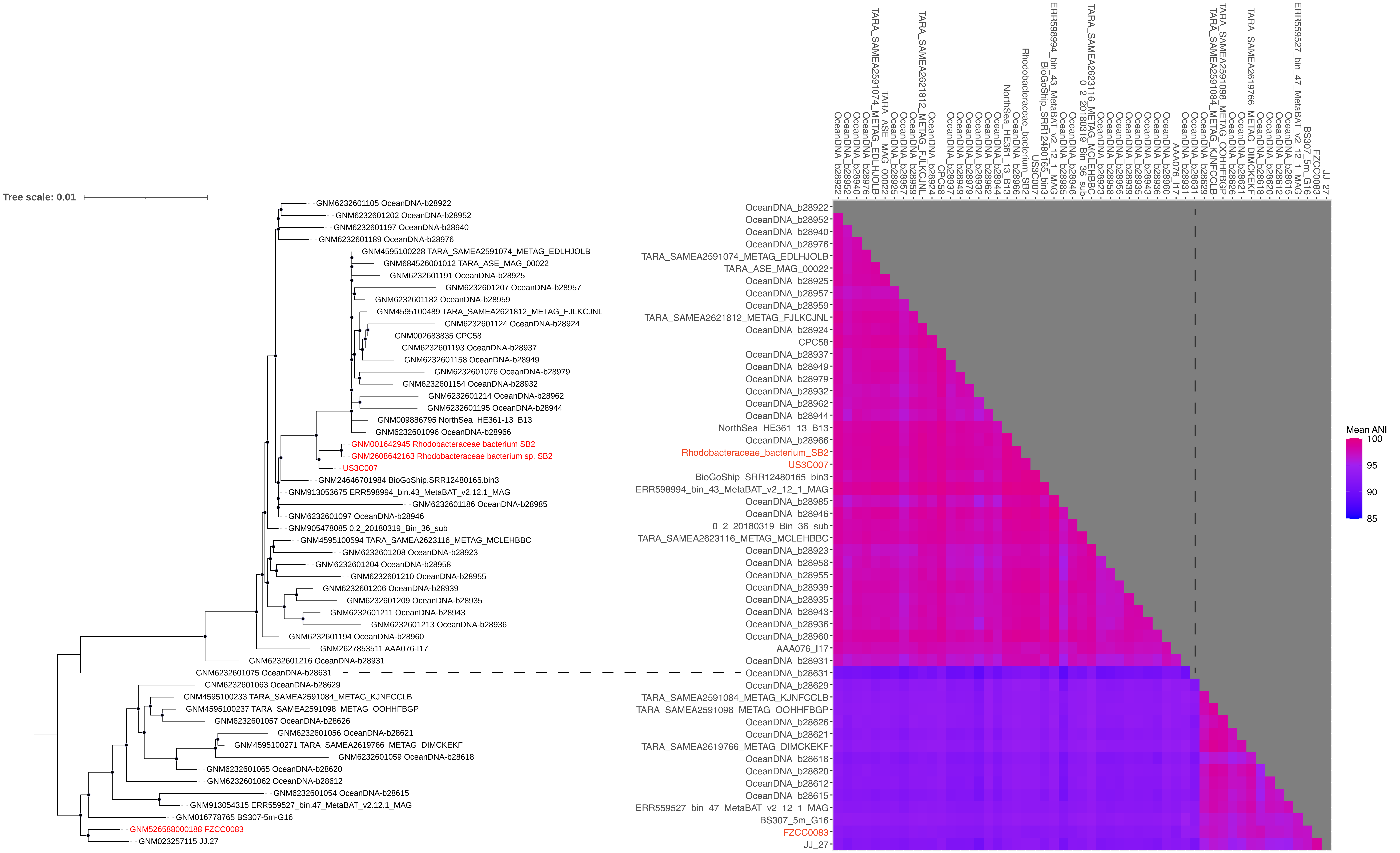

Tree scale: 0.01
GNM6232601105 OceanDNA-b28922
GNM6232601202 OceanDNA-b28952
GNM6232601197 OceanDNA-b28940
GNM6232601189 OceanDNA-b28976
GNM4595100228 TARA_SAMEA2591074_METAG_EDLHJOLB
GNM684526001012 TARA_ASE_MAG_00022
GNM6232601191 OceanDNA-b28925
GNM6232601207 OceanDNA-b28957
GNM6232601182 OceanDNA-b28959
GNM4595100489 TARA_SAMEA2621812_METAG_FJLKCJNL
GNM6232601124 OceanDNA-b28924
GNM002683835 CPC58
GNM6232601193 OceanDNA-b28937
GNM6232601158 OceanDNA-b28949
GNM6232601076 OceanDNA-b28979
GNM6232601154 OceanDNA-b28932
GNM6232601214 OceanDNA-b28962
GNM6232601195 OceanDNA-b28944
GNM009886795 NorthSea_HE361-13_B13
GNM6232601096 OceanDNA-b28966
GNM001642945 Rhodobacteraceae bacterium SB2
GNM2608642163 Rhodobacteraceae bacterium sp. SB2
US3C007
GNM24646701984 BioGoShip.SRR12480165.bin3
GNM913053675 ERR598994_bin.43_MetaBAT_v2.12.1_MAG
GNM6232601186 OceanDNA-b28985
GNM6232601097 OceanDNA-b28946
GNM905478085 0.2_20180319_Bin_36_sub
GNM4595100594 TARA_SAMEA2623116_METAG_MCLEHBBC
GNM6232601208 OceanDNA-b28923
GNM6232601204 OceanDNA-b28958
GNM6232601210 OceanDNA-b28955
GNM6232601206 OceanDNA-b28939
GNM6232601209 OceanDNA-b28935
GNM6232601211 OceanDNA-b28943
GNM6232601213 OceanDNA-b28936
GNM6232601194 OceanDNA-b28960
GNM2627853511 AAA076-I17
GNM6232601216 OceanDNA-b28931
GNM6232601075 OceanDNA-b28631
GNM6232601063 OceanDNA-b28629
GNM4595100233 TARA_SAMEA2591084_METAG_KJNFCCLB
GNM4595100237 TARA_SAMEA2591098_METAG_OOHHFBGP
GNM6232601057 OceanDNA-b28626
GNM6232601056 OceanDNA-b28621
GNM4595100271 TARA_SAMEA2619766_METAG_DIMCKEKF
GNM6232601059 OceanDNA-b28618
GNM6232601065 OceanDNA-b28620
GNM6232601062 OceanDNA-b28612
GNM6232601054 OceanDNA-b28615
GNM913054315 ERR559527_bin.47_MetaBAT_v2.12.1_MAG
GNM016778765 BS307-5m-G16
GNM526588000188 FZCC0083
GNM023257115 JJ.27
OceanDNA_b28922
OceanDNA_b28952
OceanDNA_b28940
OceanDNA_b28976
TARA_SAMEA2591074_METAG_EDLHJOLB
TARA_ASE_MAG_00022
OceanDNA_b28925
OceanDNA_b28957
OceanDNA_b28959
TARA_SAMEA2621812_METAG_FJLKCJNL
OceanDNA_b28924
CPC58
OceanDNA_b28937
OceanDNA_b28949
OceanDNA_b28979
OceanDNA_b28932
OceanDNA_b28962
OceanDNA_b28944
NorthSea_HE361_13_B13
OceanDNA_b28966
Rhodobacteraceae_bacterium_SB2
US3C007
BioGoShip_SRR12480165_bin3
ERR598994_bin_43_MetaBAT_v2_12_1_MAG
OceanDNA_b28985
OceanDNA_b28946
0_2_20180319_Bin_36_sub
TARA_SAMEA2623116_METAG_MCLEHBBC
OceanDNA_b28923
OceanDNA_b28958
OceanDNA_b28955
OceanDNA_b28939
OceanDNA_b28935
OceanDNA_b28943
OceanDNA_b28936
OceanDNA_b28960
AAA076_I17
OceanDNA_b28931
OceanDNA_b28631
OceanDNA_b28629
TARA_SAMEA2591084_METAG_KJNFCCLB
TARA_SAMEA2591098_METAG_OOHHFBGP
OceanDNA_b28626
OceanDNA_b28621
TARA_SAMEA2619766_METAG_DIMCKEKF
OceanDNA_b28618
OceanDNA_b28620
OceanDNA_b28612
OceanDNA_b28615
ERR559527_bin_47_MetaBAT_v2_12_1_MAG
BS307_5m_G16
FZCC0083
JJ_27
Mean ANI
100
95
90
85

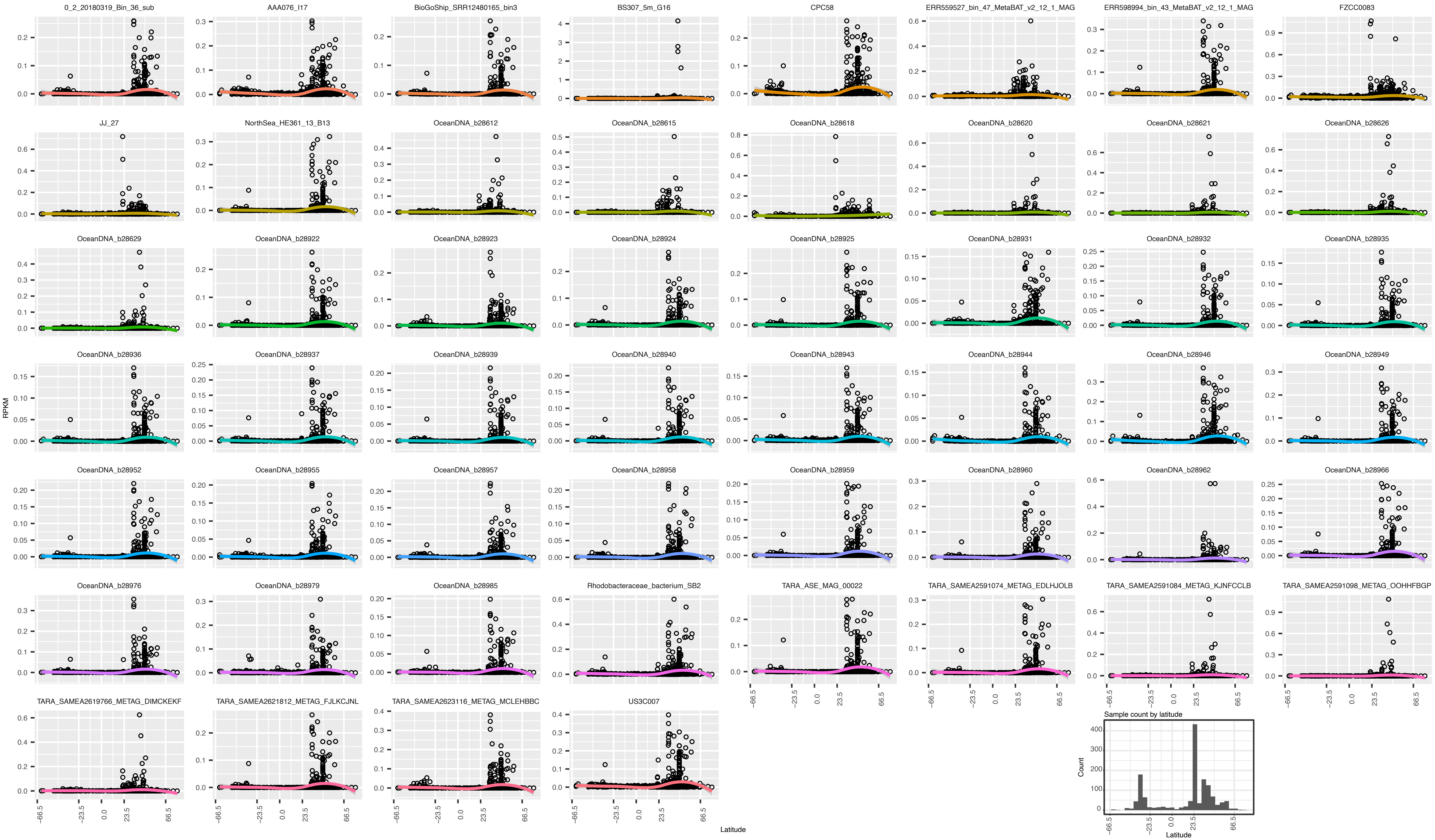
0_2_20180319_Bin_36_sub
AAA076_I17
BioGoShip_SRR12480165_bin3
BS307_5m_G16
CPC58
ERR559527_bin_47_MetaBAT_v2_12_1_MAG
ERR598994_bin_43_MetaBAT_v2_12_1_MAG
FZCC0083
JJ_27
NorthSea_HE361_13_B13
OceanDNA_b28612
OceanDNA_b28615
OceanDNA_b28618
OceanDNA_b28620
OceanDNA_b28621
OceanDNA_b28626
OceanDNA_b28629
OceanDNA_b28922
OceanDNA_b28923
OceanDNA_b28924
OceanDNA_b28925
OceanDNA_b28931
OceanDNA_b28932
OceanDNA_b28935
OceanDNA_b28936
OceanDNA_b28937
OceanDNA_b28939
OceanDNA_b28940
OceanDNA_b28943
OceanDNA_b28944
OceanDNA_b28946
OceanDNA_b28949
OceanDNA_b28952
OceanDNA_b28955
OceanDNA_b28957
OceanDNA_b28958
OceanDNA_b28959
OceanDNA_b28960
OceanDNA_b28962
OceanDNA_b28966
OceanDNA_b28976
OceanDNA_b28979
OceanDNA_b28985
Rhodobacteraceae_bacterium_SB2
TARA_ASE_MAG_00022
TARA_SAMEA2591074_METAG_EDLHJOLB
TARA_SAMEA2591084_METAG_KJNFCCLB
TARA_SAMEA2591098_METAG_OOHHFBGP
TARA_SAMEA2619766_METAG_DIMCKEKF
TARA_SAMEA2621812_METAG_FJLKCJNL
TARA_SAMEA2623116_METAG_MCLEHBBC
US3C007
RPKM
Latitude
-66.5
-23.5
0.0
23.5
66.5
Sample count by latitude
Count
0
100
200
300
400

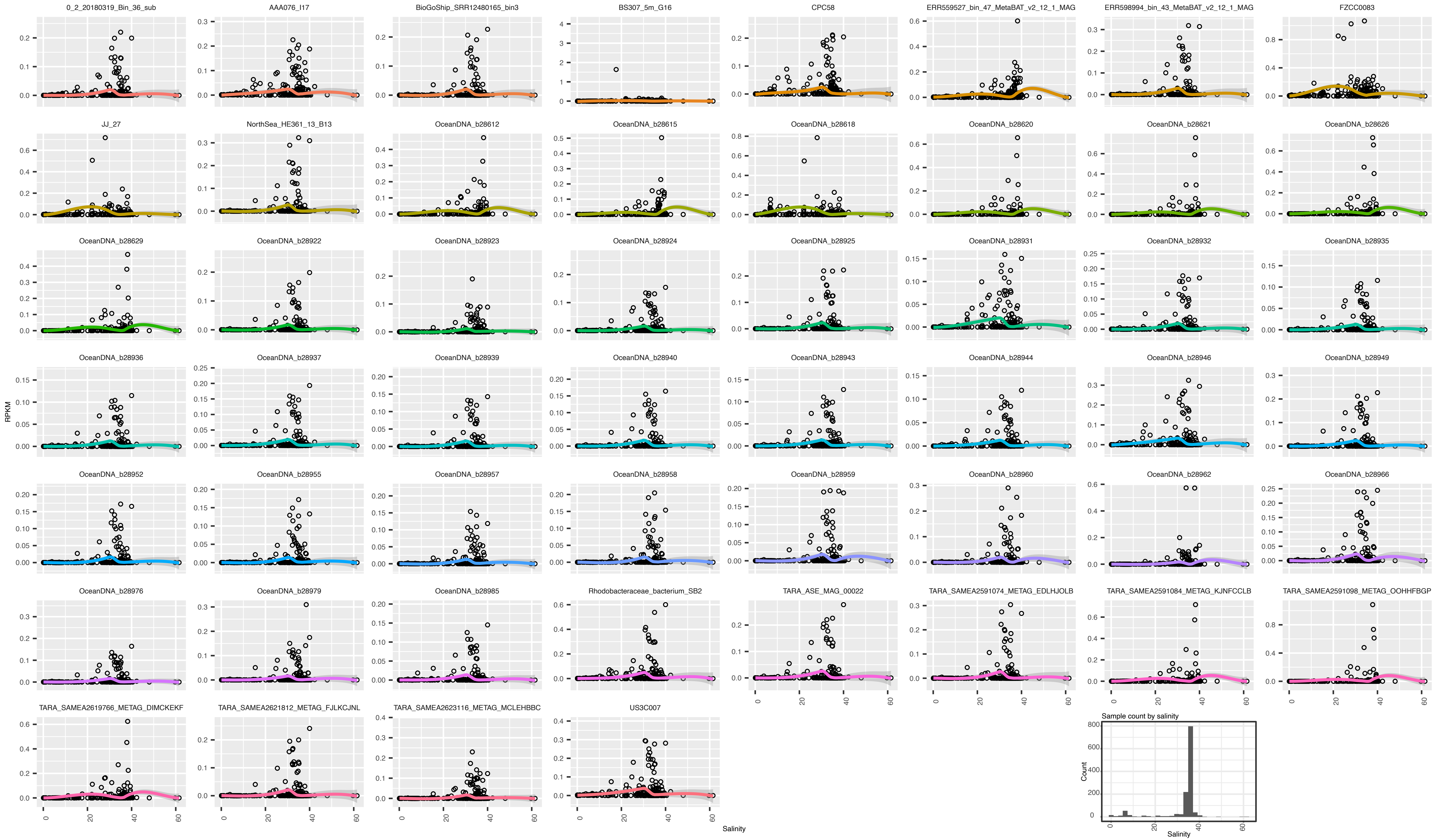

0_2_20180319_Bin_36_sub
AAA076_I17
BioGoShip_SRR12480165_bin3
BS307_5m_G16
CPC58
ERR559527_bin_47_MetaBAT_v2_12_1_MAG
ERR598994_bin_43_MetaBAT_v2_12_1_MAG
FZCC0083
JJ_27
NorthSea_HE361_13_B13
OceanDNA_b28612
OceanDNA_b28615
OceanDNA_b28618
OceanDNA_b28620
OceanDNA_b28621
OceanDNA_b28626
OceanDNA_b28629
OceanDNA_b28922
OceanDNA_b28923
OceanDNA_b28924
OceanDNA_b28925
OceanDNA_b28931
OceanDNA_b28932
OceanDNA_b28935
OceanDNA_b28936
OceanDNA_b28937
OceanDNA_b28939
OceanDNA_b28940
OceanDNA_b28943
OceanDNA_b28944
OceanDNA_b28946
OceanDNA_b28949
OceanDNA_b28952
OceanDNA_b28955
OceanDNA_b28957
OceanDNA_b28958
OceanDNA_b28959
OceanDNA_b28960
OceanDNA_b28962
OceanDNA_b28966
OceanDNA_b28976
OceanDNA_b28979
OceanDNA_b28985
Rhodobacteraceae_bacterium_SB2
TARA_ASE_MAG_00022
TARA_SAMEA2591074_METAG_EDLHJOLB
TARA_SAMEA2591084_METAG_KJNFCCLB
TARA_SAMEA2591098_METAG_OOHHFBGP
TARA_SAMEA2619766_METAG_DIMCKEKF
TARA_SAMEA2621812_METAG_FJLKCJNL
TARA_SAMEA2623116_METAG_MCLEHBBC
US3C007
RPKM
Salinity
Sample count by salinity
Count
Salinity

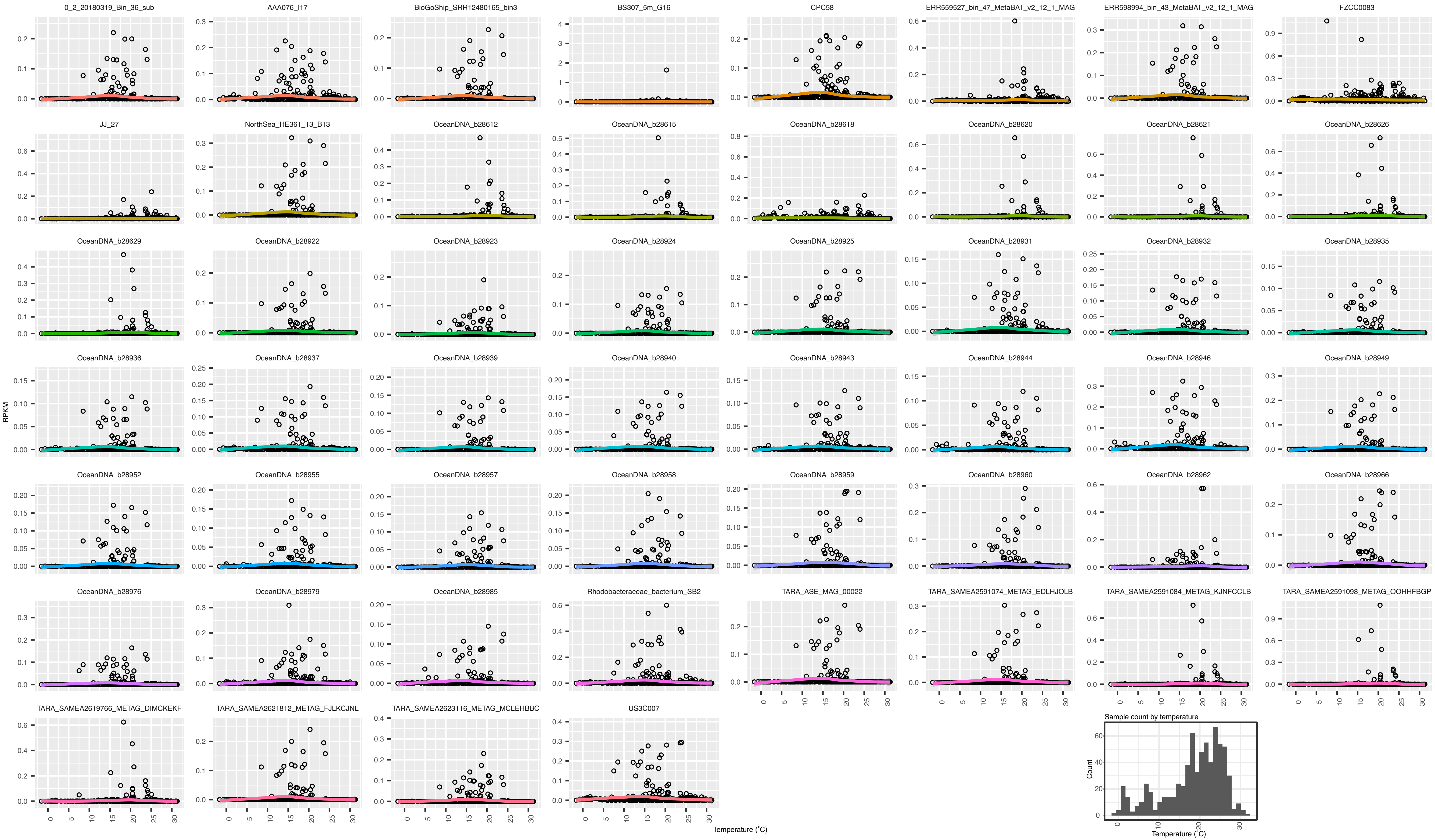
0_2_20180319_Bin_36_sub
AAA076_I17
BioGoShip_SRR12480165_bin3
BS307_5m_G16
CPC58
ERR559527_bin_47_MetaBAT_v2_12_1_MAG
ERR598994_bin_43_MetaBAT_v2_12_1_MAG
FZCC0083
JJ_27
NorthSea_HE361_13_B13
OceanDNA_b28612
OceanDNA_b28615
OceanDNA_b28618
OceanDNA_b28620
OceanDNA_b28621
OceanDNA_b28626
OceanDNA_b28629
OceanDNA_b28922
OceanDNA_b28923
OceanDNA_b28924
OceanDNA_b28925
OceanDNA_b28931
OceanDNA_b28932
OceanDNA_b28935
OceanDNA_b28936
OceanDNA_b28937
OceanDNA_b28939
OceanDNA_b28940
OceanDNA_b28943
OceanDNA_b28944
OceanDNA_b28946
OceanDNA_b28949
OceanDNA_b28952
OceanDNA_b28955
OceanDNA_b28957
OceanDNA_b28958
OceanDNA_b28959
OceanDNA_b28960
OceanDNA_b28962
OceanDNA_b28966
OceanDNA_b28976
OceanDNA_b28979
OceanDNA_b28985
Rhodobacteraceae_bacterium_SB2
TARA_ASE_MAG_00022
TARA_SAMEA2591074_METAG_EDLHJOLB
TARA_SAMEA2591084_METAG_KJNFCCLB
TARA_SAMEA2591098_METAG_OOHHFBGP
TARA_SAMEA2619766_METAG_DIMCKEKF
TARA_SAMEA2621812_METAG_FJLKCJNL
TARA_SAMEA2623116_METAG_MCLEHBBC
US3C007
RPKM
Temperature (°C)
Sample count by temperature
Count
Temperature (°C)

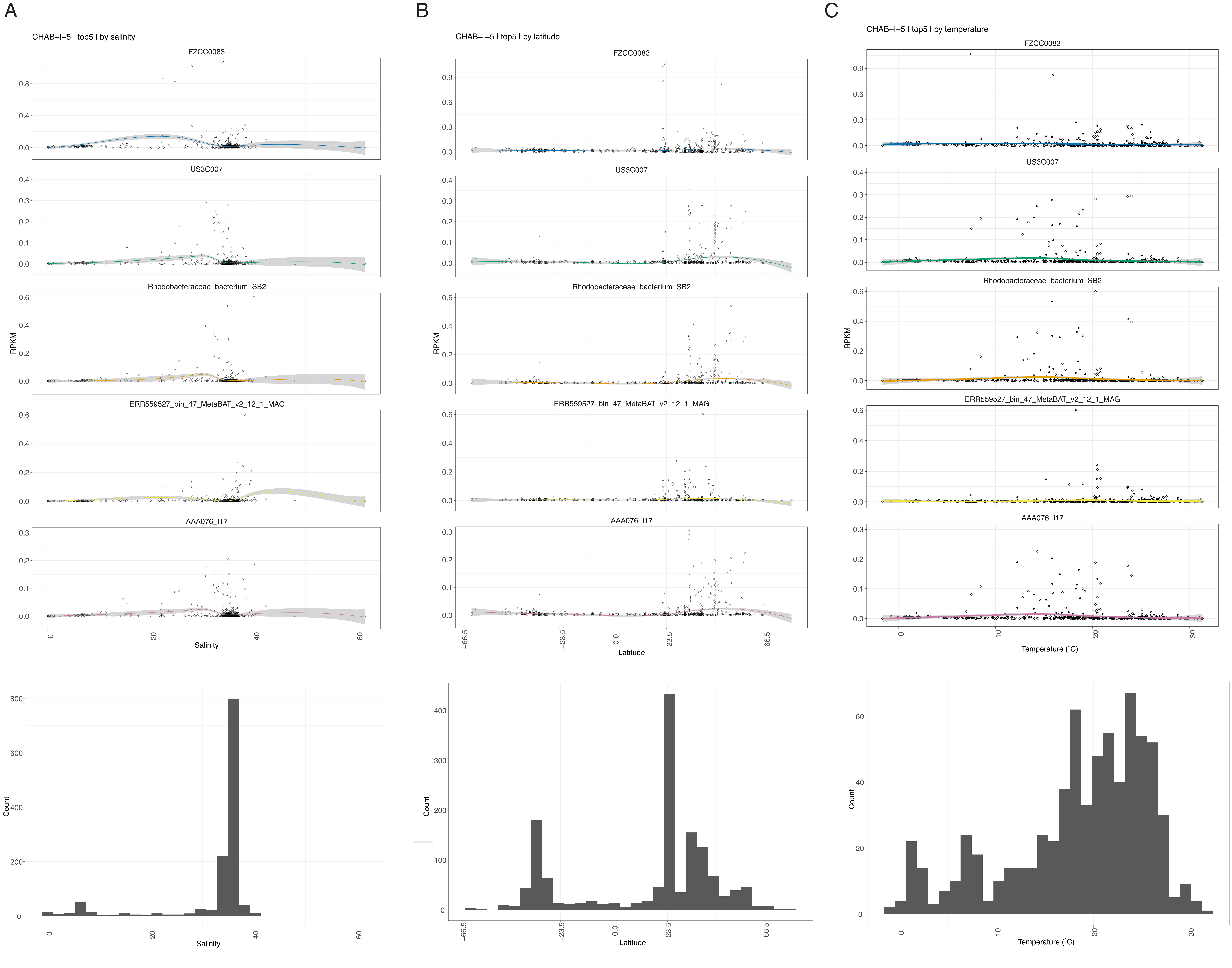
A
B
C
CHAB-I-5 | top5 | by salinity
CHAB-I-5 | top5 | by latitude
CHAB-I-5 | top5 | by temperature
FZCC0083
US3C007
Rhodobacteraceae_bacterium_SB2
ERR559527_bin_47_MetaBAT_v2_12_1_MAG
AAA076_I17
RPKM
Salinity
Latitude
Temperature (°C)
Count

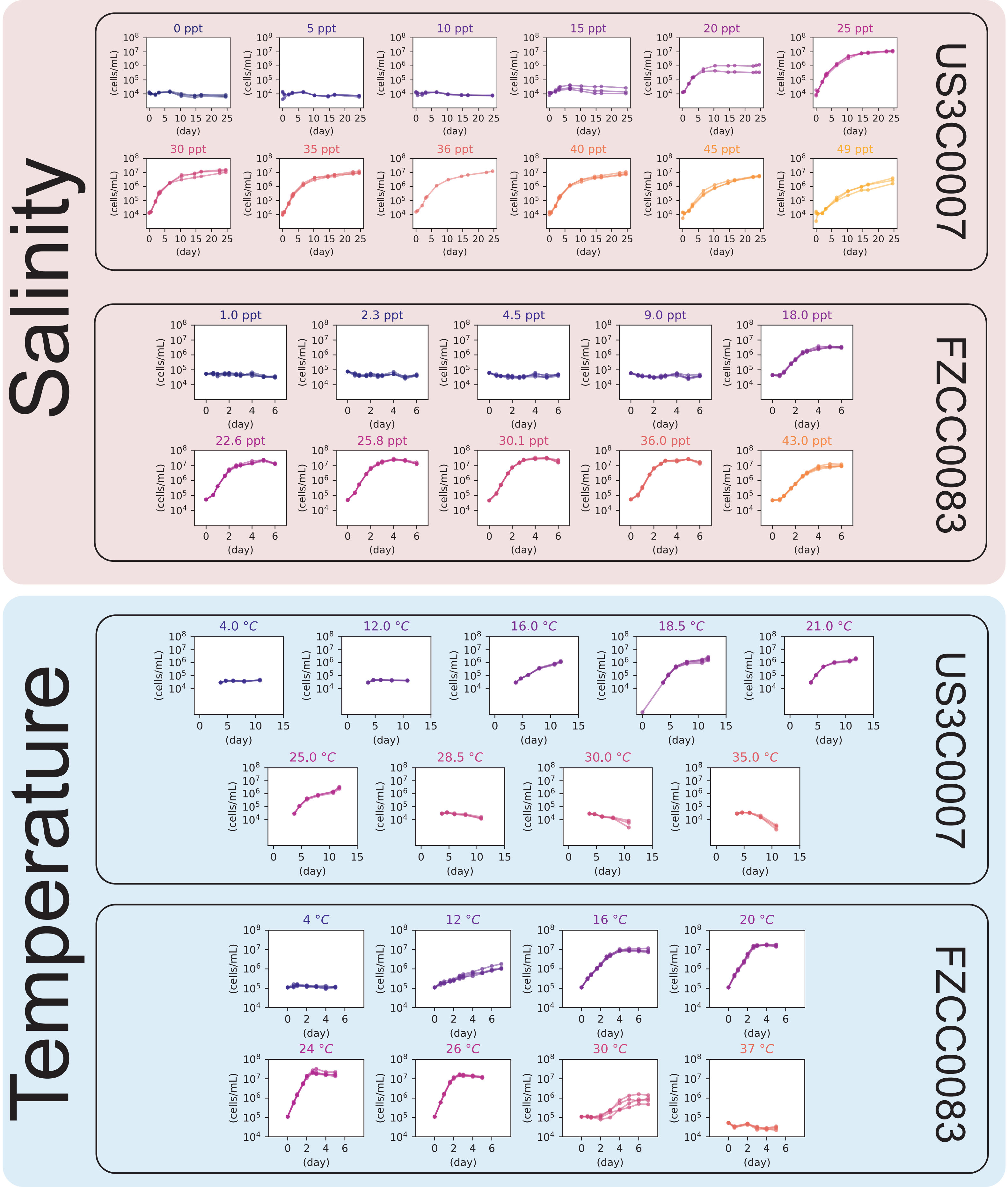

Salinity
US3C0007
0 ppt
5 ppt
10 ppt
15 ppt
20 ppt
25 ppt
30 ppt
35 ppt
36 ppt
40 ppt
45 ppt
49 ppt
FZCC0083
1.0 ppt
2.3 ppt
4.5 ppt
9.0 ppt
18.0 ppt
22.6 ppt
25.8 ppt
30.1 ppt
36.0 ppt
43.0 ppt
Temperature
US3C0007
4.0 °C
12.0 °C
16.0 °C
18.5 °C
21.0 °C
25.0 °C
28.5 °C
30.0 °C
35.0 °C
FZCC0083
4 °C
12 °C
16 °C
20 °C
24 °C
26 °C
30 °C
37 °C
(cells/mL)
(day)

$$V = \frac{4}{3}\pi r^3 + \pi r^2 h$$

$$S = \pi r^2 + 2rh$$

$$h = (S - \pi r^2)/2r$$

$$l = h + 2r$$

$$V = \frac{4}{3}\pi r^3 + \pi r^2 \cdot (S - \pi r^2)/2r$$

$$= \frac{4}{3}\pi r^3 + \pi r \cdot (S - \pi r^2)$$

$$= \frac{4}{3}\pi r^3 + S \cdot \pi r - \pi \cdot \pi r^3$$

$$= (\frac{4}{3} - \pi)\pi r^3 + S \cdot \pi r$$

$$SA = 2\pi r(2r + h)$$

$$= 2\pi r(2r + (S - \pi r^2)/2r)$$

$$= 4\pi r^2 + \frac{2\pi r(S - \pi r^2)}{2r}$$

$$= 4\pi r^2 + \pi S - \pi^2 r^2$$

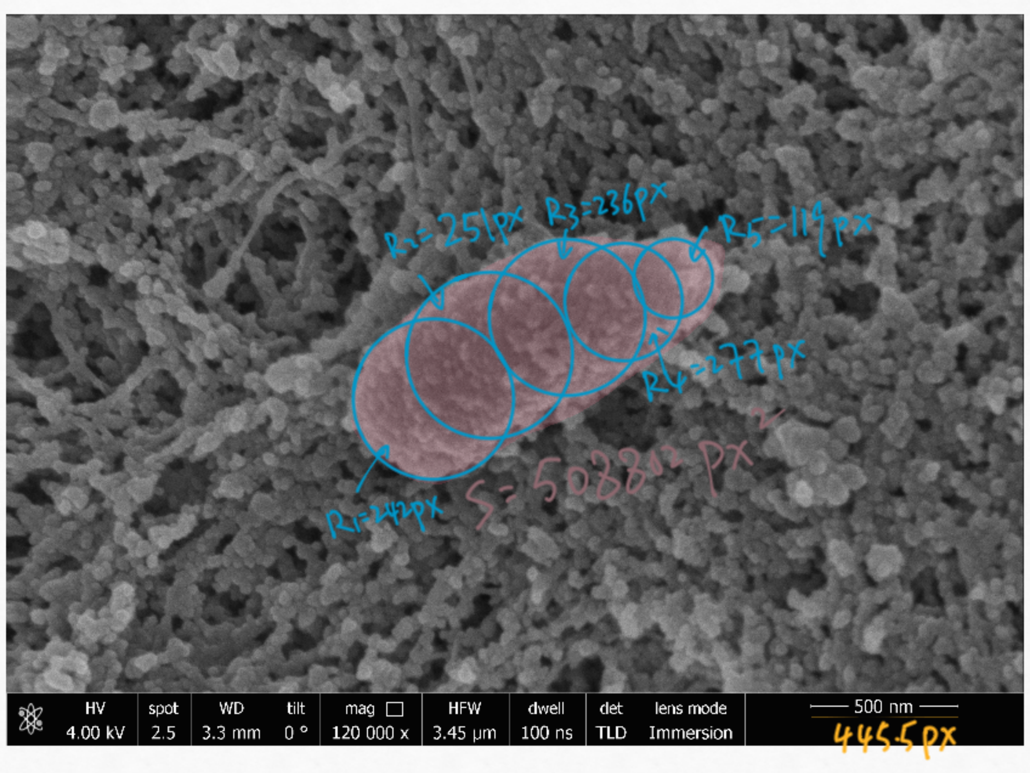
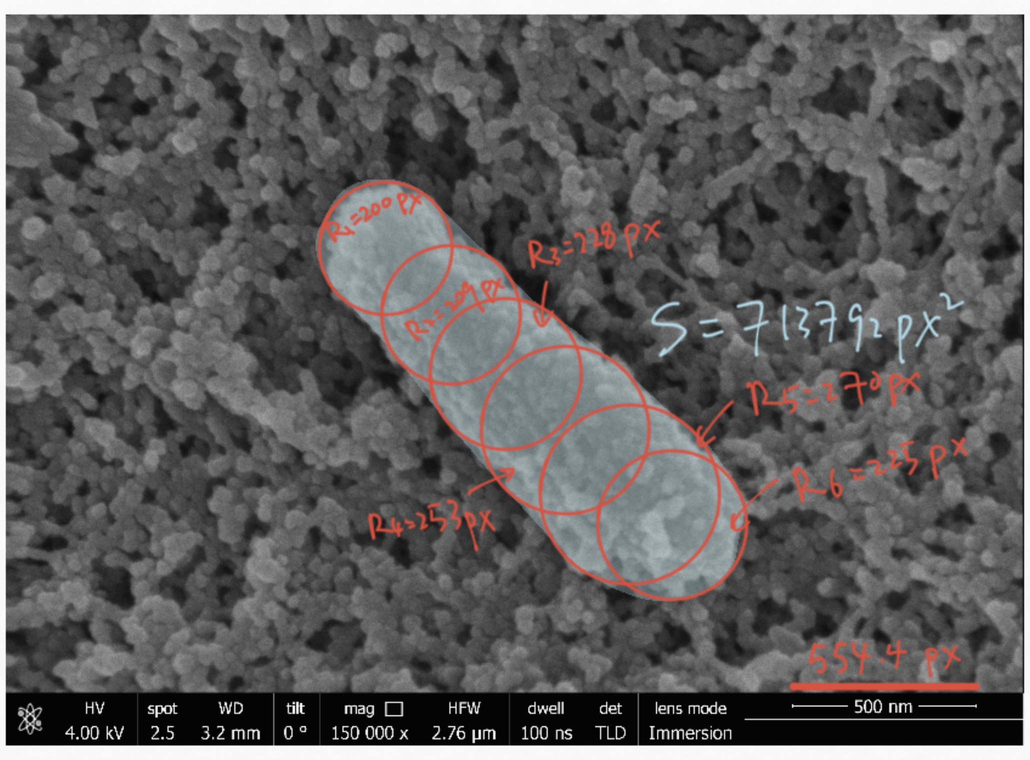
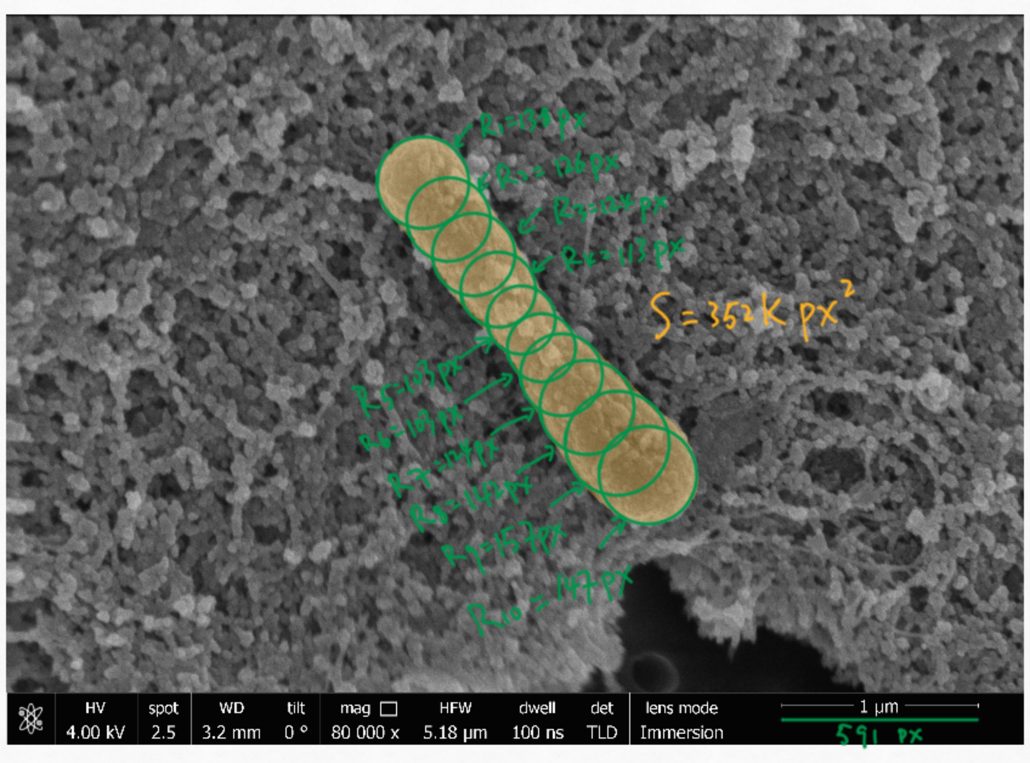
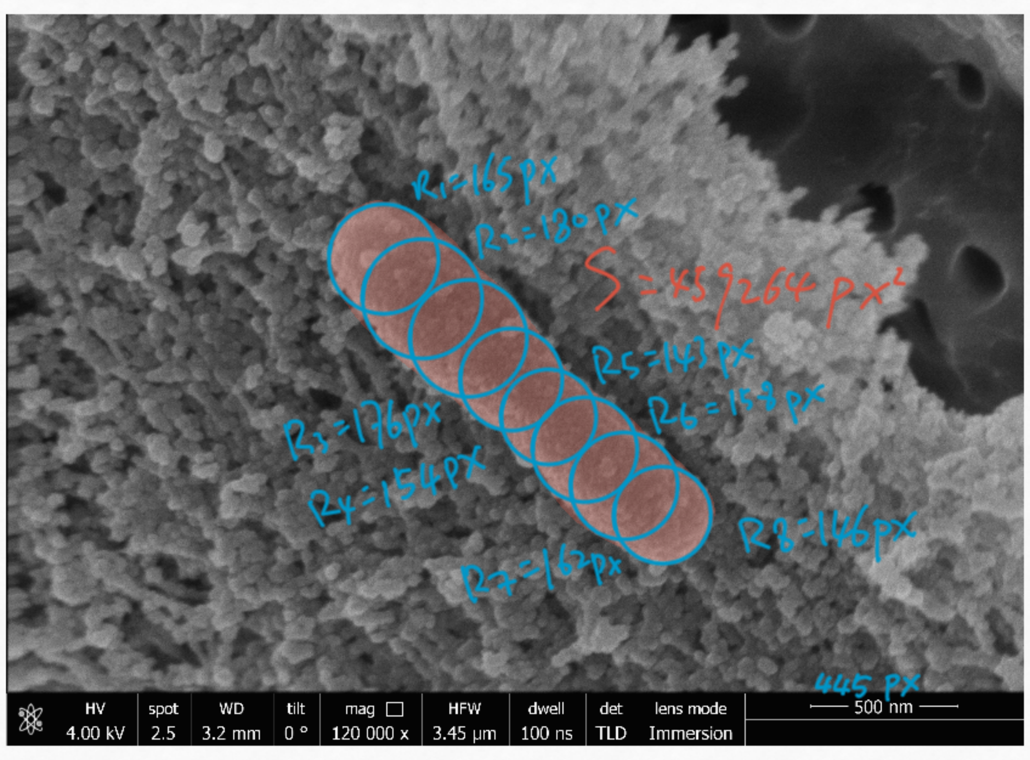
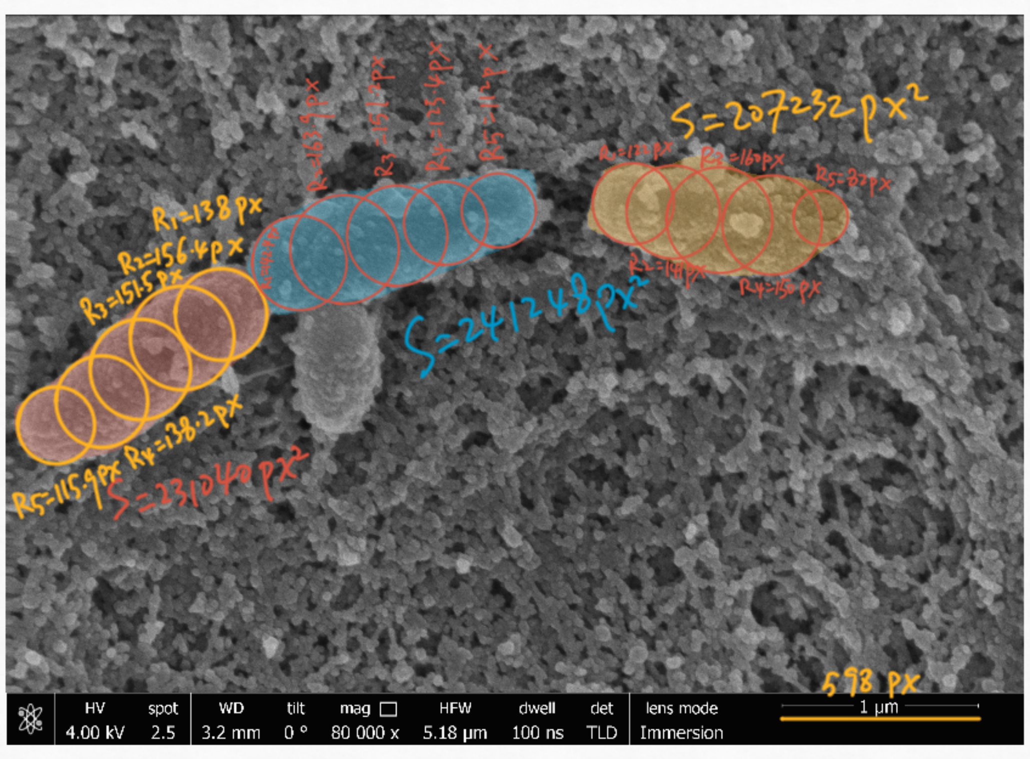
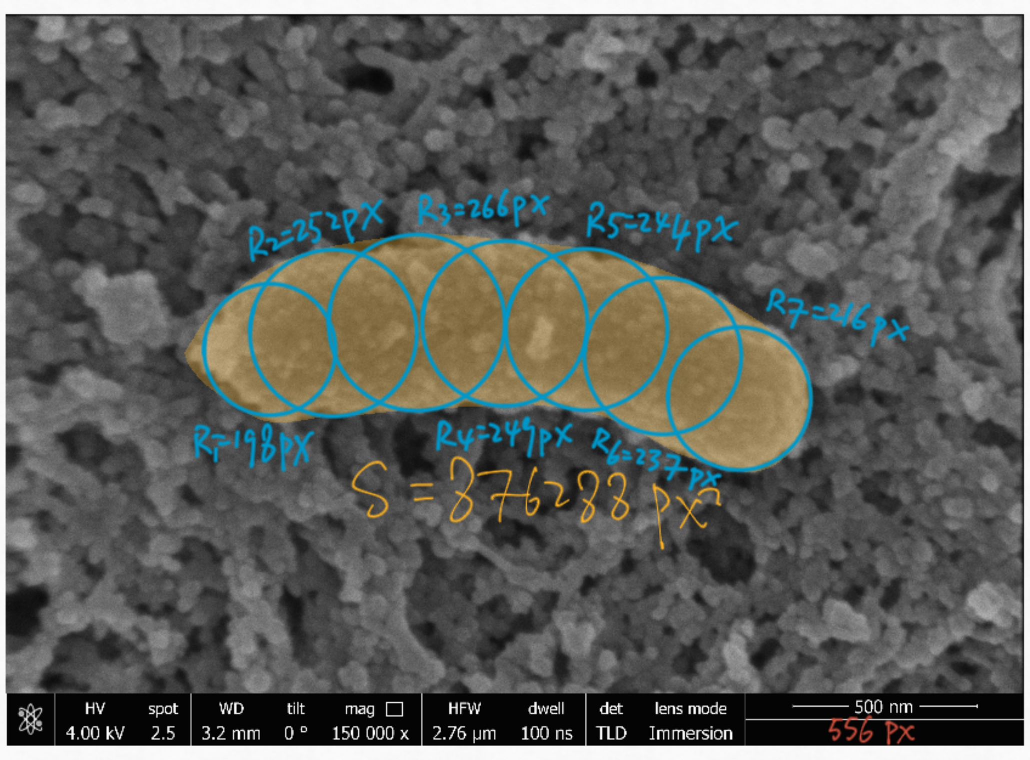
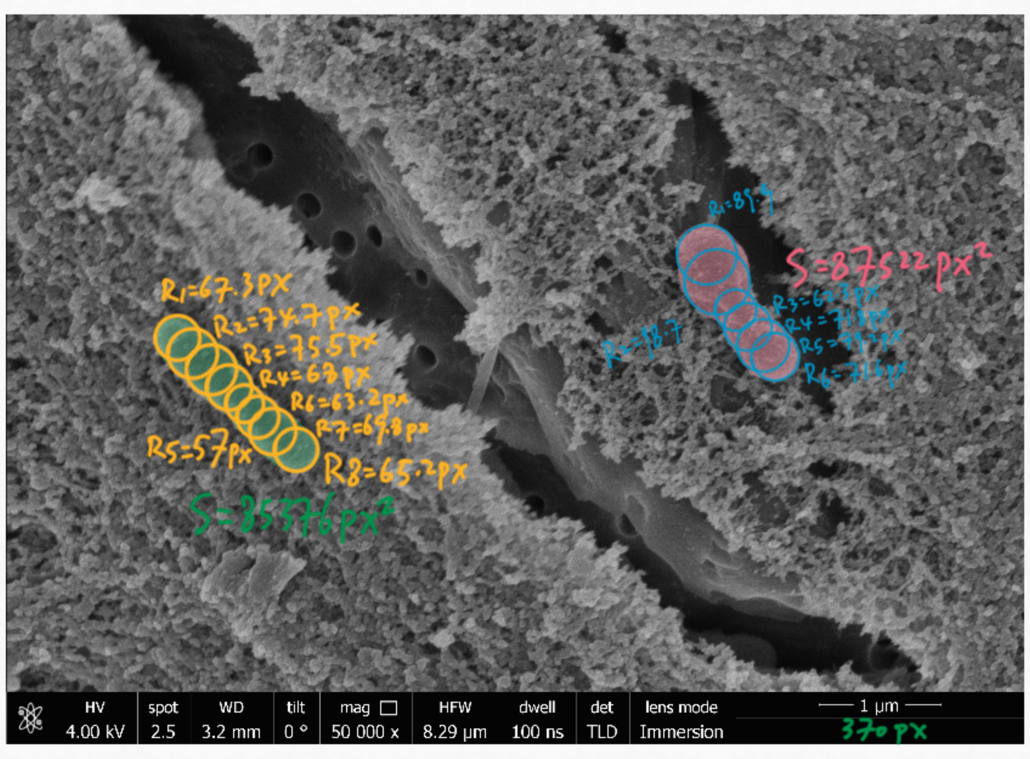
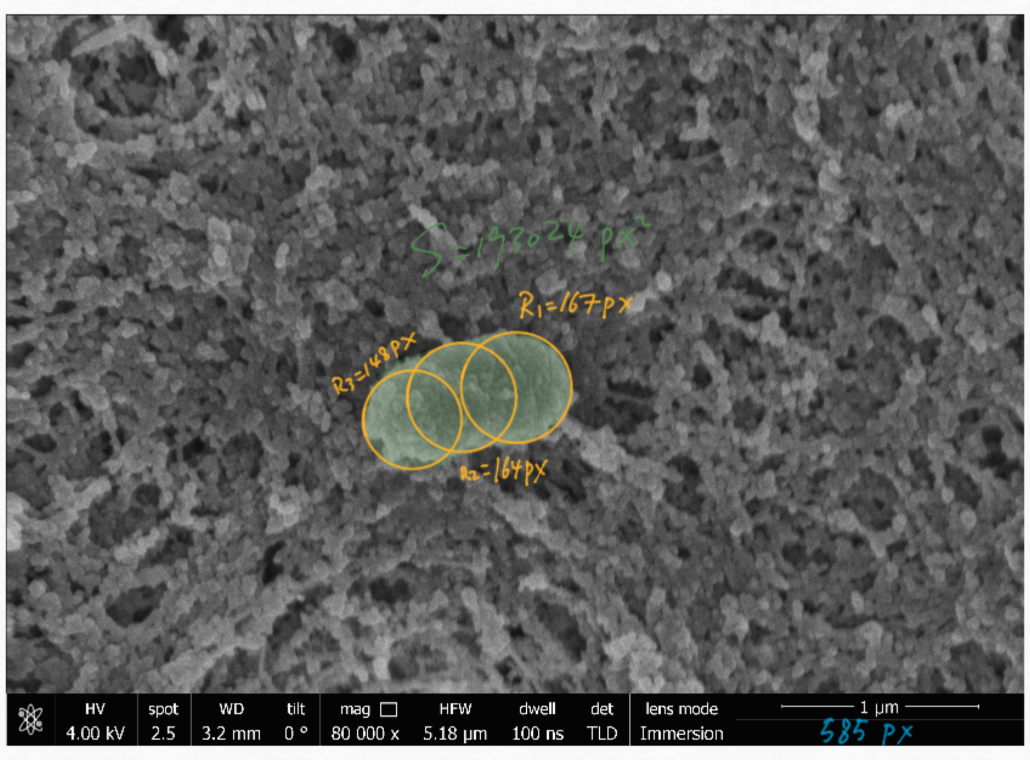
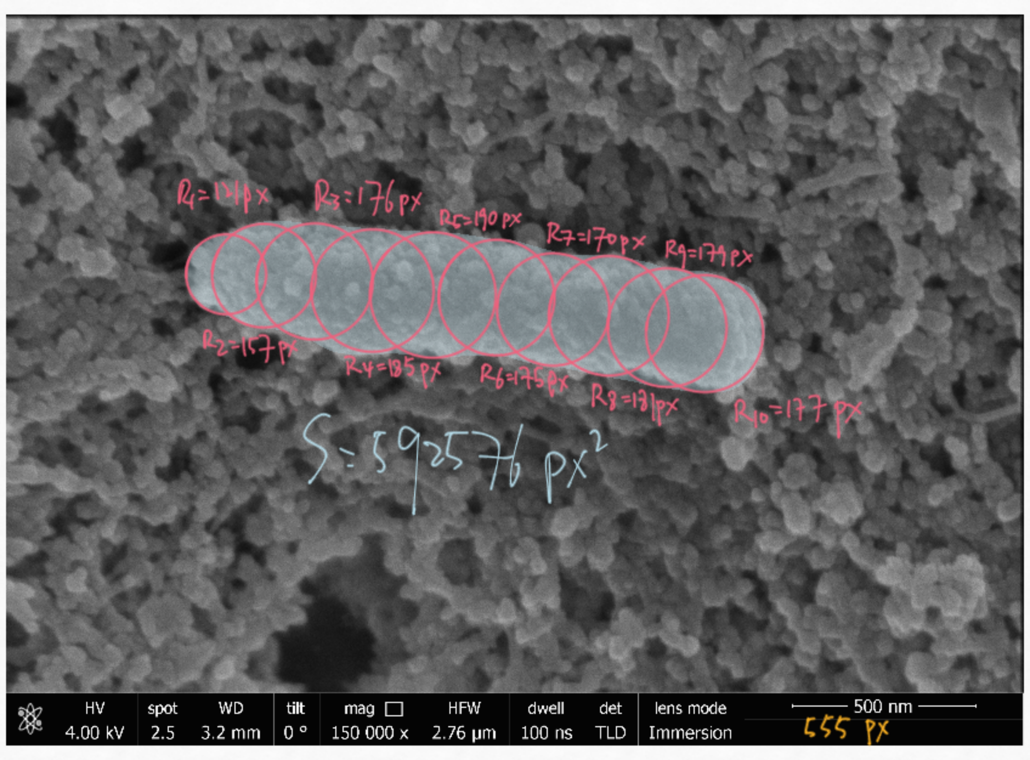
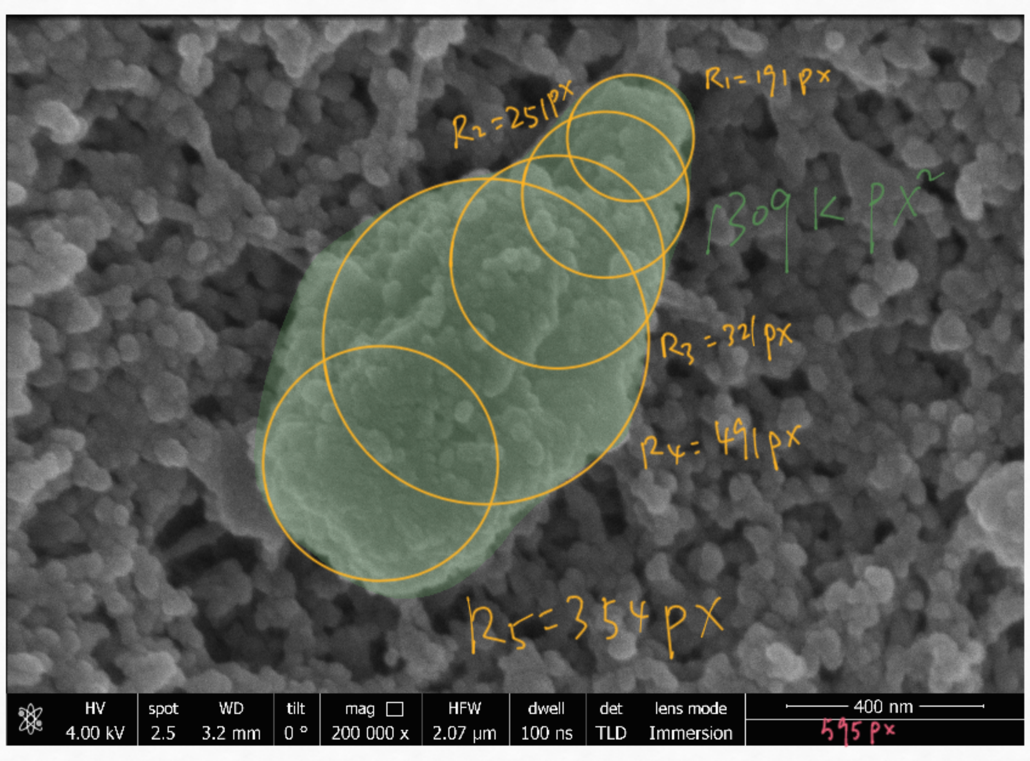
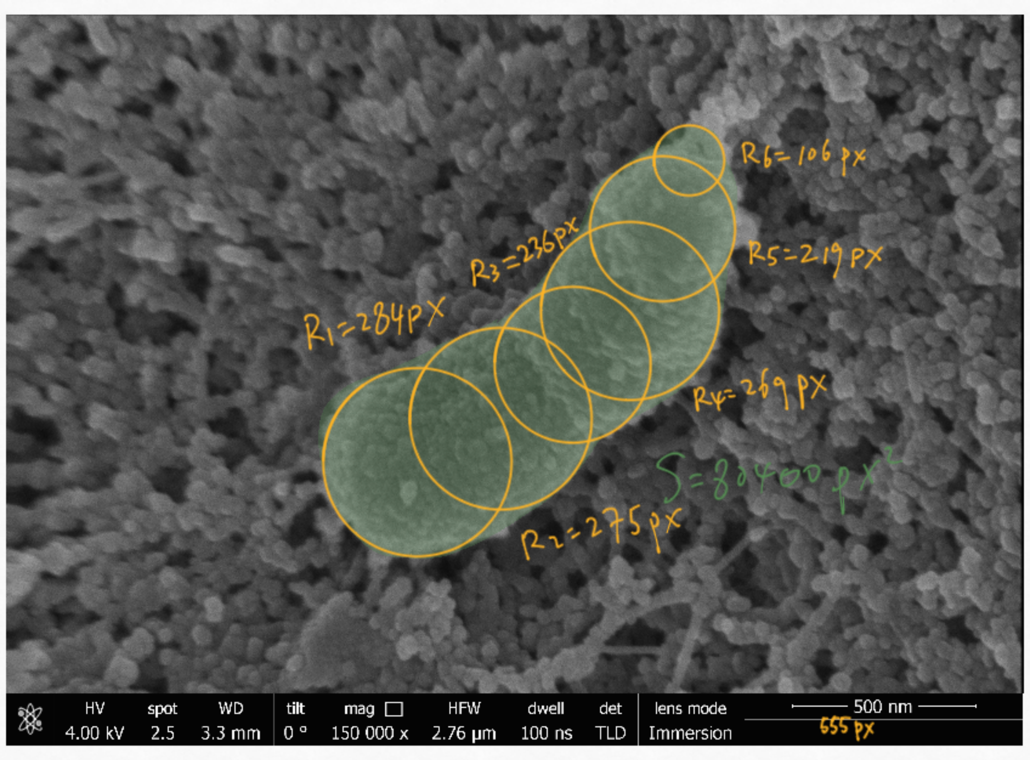
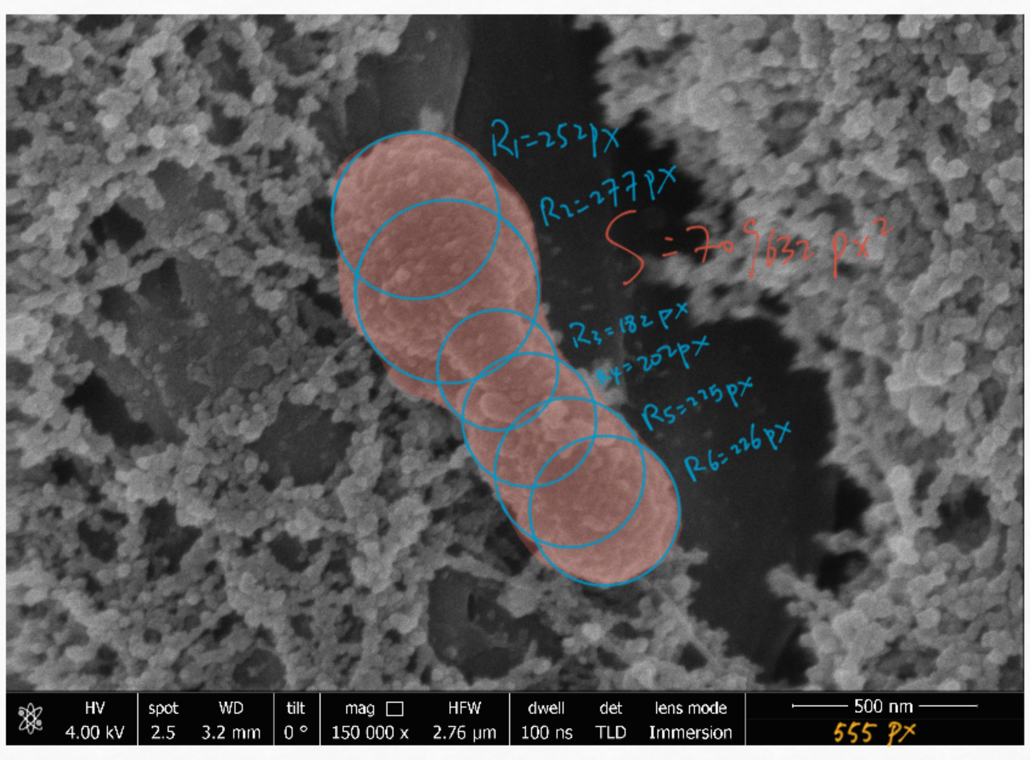
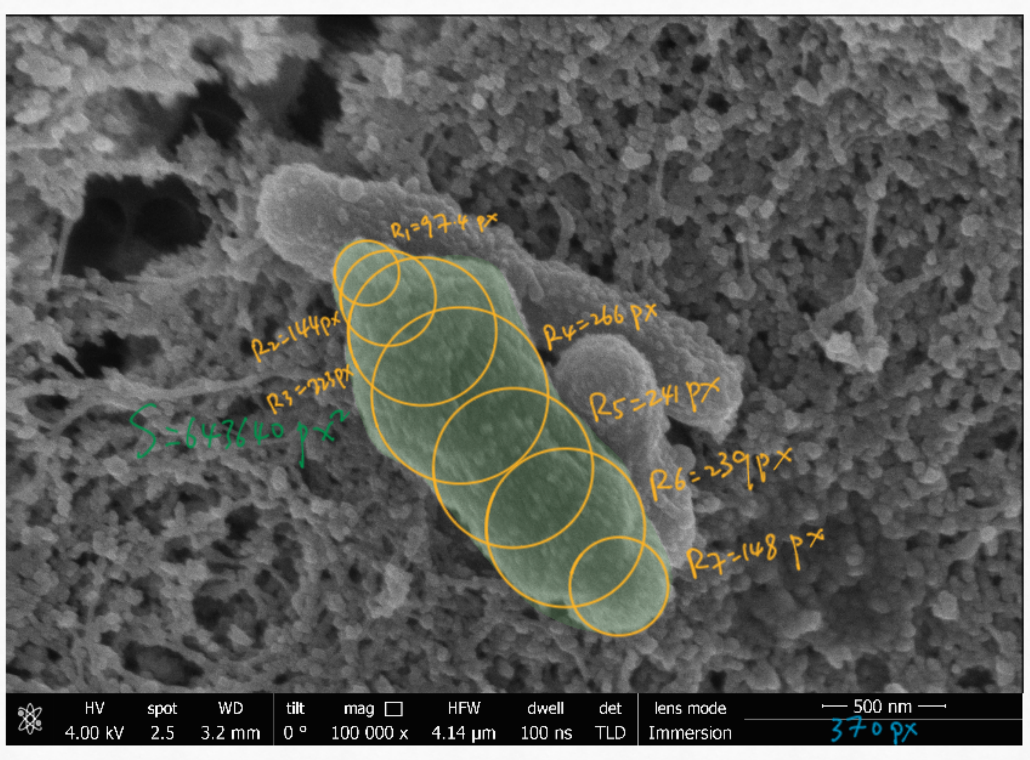
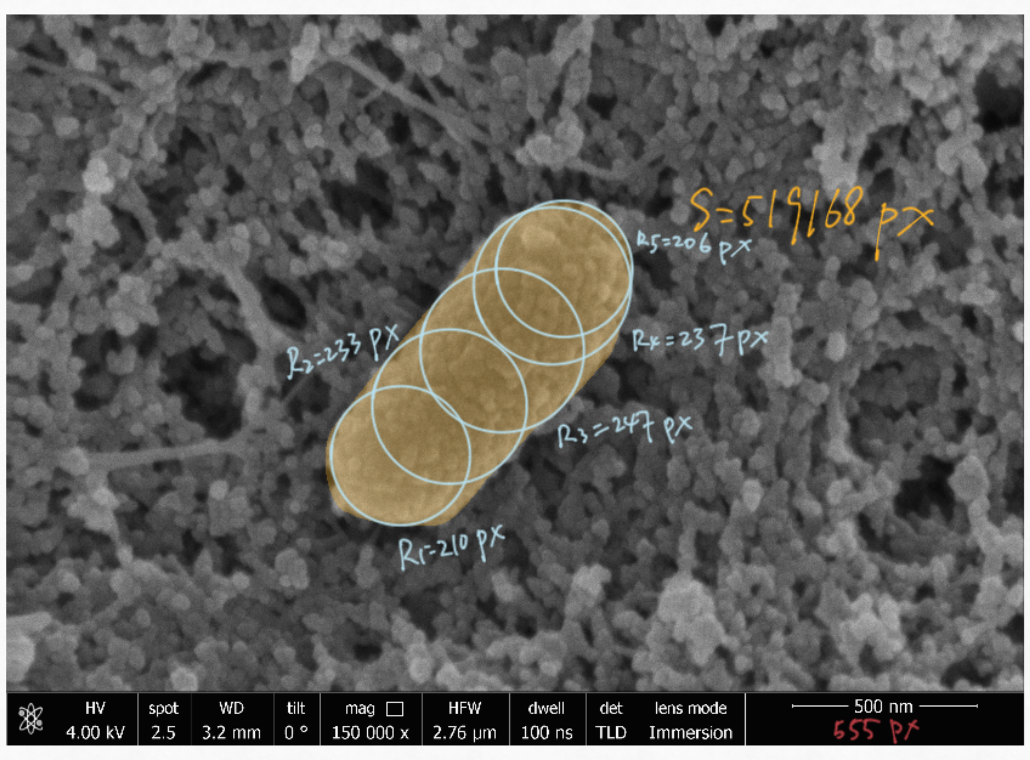
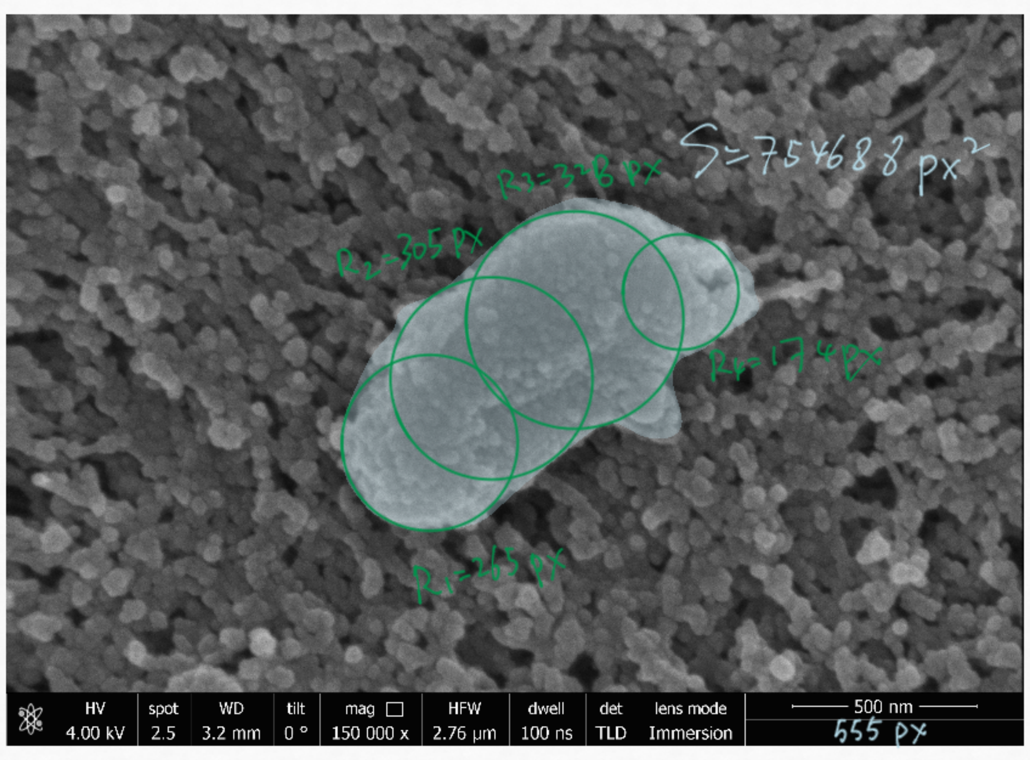

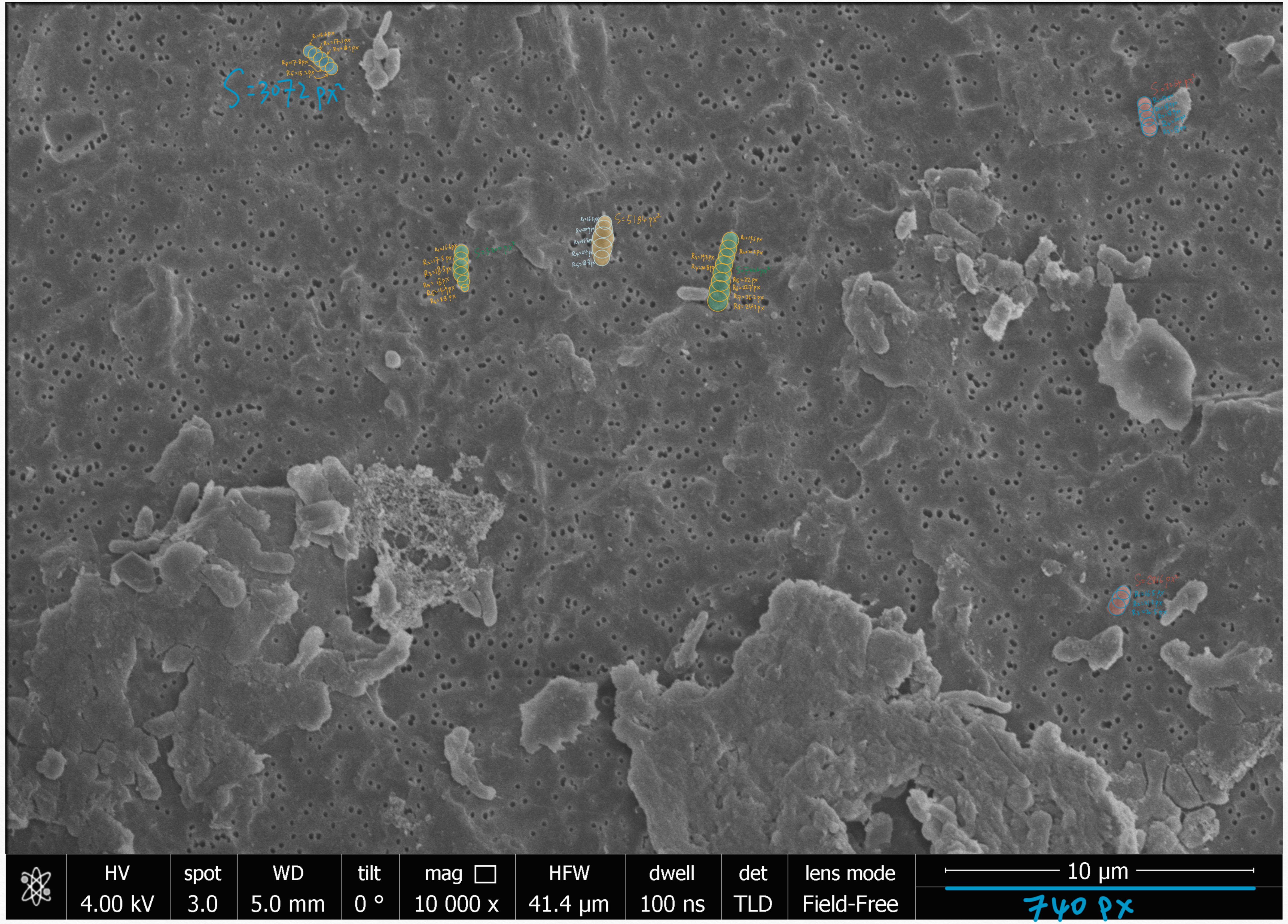

HV
4.00 kV
spot
3.0
WD
5.0 mm
tilt
0 °
mag
10 000 x
HFW
41.4 µm
dwell
100 ns
det
TLD
lens mode
Field-Free
10 µm
740 px


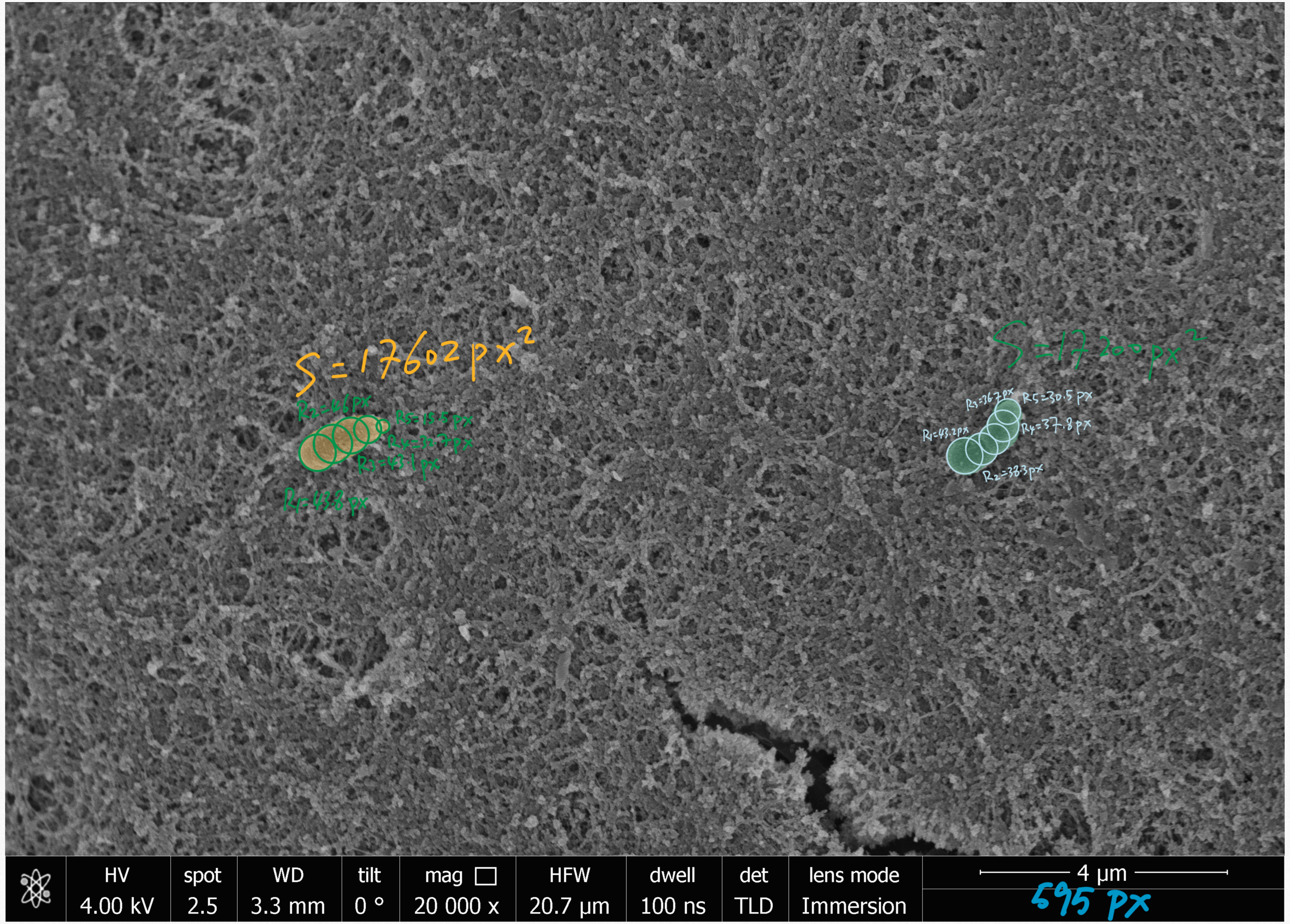

S=17602 px²
HV
4.00 kV
spot
2.5
WD
3.3 mm
tilt
0 °
mag
20 000 x
HFW
20.7 µm
dwell
100 ns
det
TLD
lens mode
Immersion
4 µm
595 px

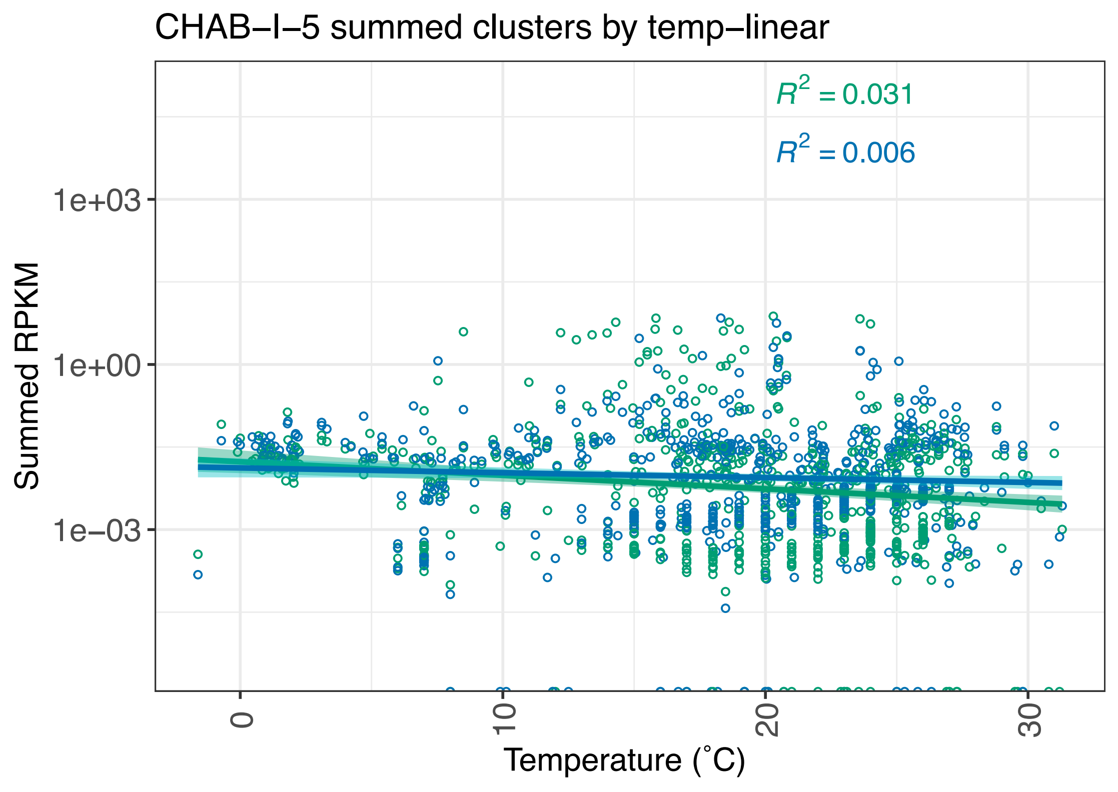

CHAB–I–5 summed clusters by temp–linear
$R^2 = 0.031$
$R^2 = 0.006$
1e+03
1e+00
1e–03
Summed RPKM
0
10
20
30
Temperature (˚C)
